# Comparative Genomic and Functional Profiling of ECM-Targeting Enzymes in *Bacteroides*, a Key Genus of the Human Gut Microbiome

**DOI:** 10.64898/2026.03.04.709643

**Authors:** Karen M Mancera Azamar, Krishna Rajesh, Bryanna Downing, Marcos Javith, Isabela Yamhure, Ana Maria Porras

## Abstract

Purpose

The human extracellular matrix (ECM) provides essential cues for intestinal homeostasis. While most studies focus on ECM degradation by host cells, our prior work suggests that commensal gut microbes may also contribute to these remodeling processes. Here, we continue exploring this novel dimension of host-microbe interactions by profiling the proteolytic diversity and substrate-specific activity of ECM-targeting enzymes across species of *Bacteroides,* a dominant and metabolically versatile gut genus.

**Methods:** We curated a custom ECM-specific enzyme database from the BRENDA repository and used it to perform comparative genomic analyses across 11 *Bacteroides* species, mapping the diversity and abundance of candidate ECM-degrading proteases and carbohydrate active enzymes (CAZymes). Functional activity was evaluated via *in vitro* degradation assays using purified substrates. Family-specific protease inhibitors were used to confirm the major catalytic classes involved.

**Results:** ECM-targeting CAZymes and proteases were broadly encoded across all 11 genomes, with gene counts positively correlated with genome size and GAG-associated genes comprising the largest substrate category. Experimental degradation assays revealed species- and substrate-specific activity patterns, including elastin degradation restricted to a subset of species, a capacity previously undocumented in intestinal *Bacteroides*. Genomic predictions showed limited concordance with measured enzymatic activity, suggesting context-dependent regulation of ECM-degrading enzymes. Inhibitor experiments confirmed that collagen degradation is driven primarily by metalloproteases and secondarily by serine proteases across representative species.

**Conclusions:** Our findings position commensal *Bacteroides* as a rich, yet underappreciated, source of ECM-degrading enzymes. This work underscores the need to consider microbiota as key modulators of host tissue homeostasis and potential targets for therapeutic modulation.

**BIOGRAPHY:** Dr. Ana Maria Porras is an Assistant Professor of Biomedical Engineering at the University of Florida, where she leads the Tissue-Microbe Interactions lab. Her group leverages cell and tissue engineering, bioinformatics, and statistical modeling to understand how microorganisms regulate human extracellular matrix remodeling. Her work centers primarily on the gut microbiome, cardiovascular health, and tropical infectious diseases. Dr. Porras is also a science artist, and a science communicator, particularly in interested in evidence-based, culturally informed, and multilingual practices to improve public engagement with science. She is the co-founder and Senior Advisor of the Latinx in Biomedical Engineering community, and the recipient of multiple awards, including the UF Excellence Award for Assistant Professors, the NSF Faculty Early Career Development (CAREER) Award, the NIH Maximizing Investigators Research Award (MIRA), the AAAS Early Career Award for Public Engagement with Science, and and the Rising Star Award from the Academy of Science, Engineering, and Medicine of Florida. Prior to arriving in Florida, Dr. Porras was a Presidential Postdoctoral Fellow at Cornell University. She holds a B.S. in biomedical engineering from the University of Texas at Austin, and a Ph.D. from the University of Wisconsin-Madison, where she was an American Heart Association Predoctoral Fellow.

## INTRODUCTION

The intestinal extracellular matrix (ECM) is a dynamic structure essential for tissue homeostasis, providing biochemical and mechanical cues that regulate epithelial barrier function, immune surveillance, and wound repair [1–3]. This complex network comprises structural proteins such as collagens, elastin, and fibronectin, as well as proteoglycans and glycosaminoglycans that modulate growth factor availability and cellular signaling [4]. Intestinal ECM remodeling, the coordinated degradation and synthesis of these components, occurs in proximity to one of the densest and most metabolically active microbial communities in the human body: the gut microbiome [5]. Despite this spatial and biological overlap, the potential contribution of microbial enzymatic activity to intestinal ECM remodeling remains poorly characterized.

ECM degradation has been extensively characterized in the context of host cell biology [6, 7]. Fibroblasts, immune cells, and epithelial cells secrete a well-studied arsenal of proteases, including matrix metalloproteinases (MMPs) and cathepsins, as well as glycosidases such as hyaluronidases and heparanases [8–10]. These enzymes regulate ECM remodeling during development, wound healing, and disease progression, with their dysregulation implicated in conditions ranging from fibrosis to cancer [11–17]. Given the extensive body of work on host-derived enzymes, ECM-specific proteolytic or glycosidic activity detected in intestinal tissues, luminal samples, or stool is commonly attributed to host cells as the primary source [18–20].

Yet, gut bacteria are known to secrete diverse enzymes involved in nutrient acquisition and niche adaptation, including both proteases and carbohydrate-active enzymes (CAZymes) [21–23]. While proteolytic activity is abundant in fecal and intestinal samples rich in microbiota [24–27], the relative contributions of microbial versus host enzymes are rarely distinguished. Growing evidence challenges the assumption that ECM degradation is exclusively mediated by the host. Bacterial pathogens have long been known to bind and degrade ECM components as a mechanism of tissue invasion [28–30]. For example, in the oral cavity, bacterial trypsin-like peptidases secreted by periodontal bacteria degrade proteins in the basal lamina, contributing to loss of gingival attachment to the tooth [28, 29]. Extending this to the gut, prior work from our group demonstrated that multiple commensal bacteria species can directly degrade collagens, glycoproteins, and glycosaminoglycans *in vitro* through secreted metalloproteases, serine proteases, and a wide variety of CAZymes [31]. However, the diversity, substrate specificity, and enzymatic mechanisms underlying microbial ECM degradation in the intestine remain poorly characterized.

*Bacteroides* species are dominant members of the human gut microbiome and are renowned for their extensive CAZyme repertoires and proteolytic capacity, making them efficient degraders of complex host-derived substrates including mucins and dietary polysaccharides [32–35]. Notably, several species have been associated with intestinal diseases involving aberrant ECM remodeling, including inflammatory bowel disease and colorectal cancer [36–38]. Additionally, specific examples of ECM-targeting activity have been documented: enterotoxigenic *B. fragilis* strains secrete zinc metalloproteases capable of degrading gelatin and collagen [39, 40], while *B. thetaiotaomicron* produces sulfatases that cleave chondroitin sulfate and heparan sulfate [41–43]. Consistent with these observations, *Bacteroides* species exhibited the highest proteolytic activity against ECM substrates among the gut commensals tested in our prior work [31]. Despite this evidence, systematic profiling of ECM-targeting enzymes across the *Bacteroides* genus remains limited.

Here, we combine comparative genomics with functional assays to profile the diversity and substrate specificity of ECM-degrading proteases and CAZymes across 11 *Bacteroides* species. The panel of species selected in this study encompasses well-characterized species exhibiting prominent enzymatic functions, such as the aforementioned *B. fragilis* and *B. thetaiotaomicron*, but it also includes lesser-known species important to gut homeostasis whose enzymatic potential are less defined. For instance, *B. caccae* and *B. acidifaciens* have been observed to fluctuate in abundance in the context of mucosal inflammation [44], and disease-associated dysbiosis [45, 46]. Other species like *B. salyersiae* or *B. xylanisolvens* are garnering attention for their emerging roles in glycosaminoglycan degradation [47, 48]. Our work reveals conserved and species-specific enzymatic patterns, shedding light on potential mechanisms by which these key gut microbes interact with and modulate the intestinal extracellular matrix.

## RESULTS

### ECM-specific CAZymes and proteases are widely encoded across *Bacteroides* genomes and increase with genome size

We began our analysis by evaluating the genomic distribution of ECM-targeting enzymes across the genus, we analyzed 11 publicly available *Bacteroides* genomes representing species commonly present in the human gut (**Table 1**). To characterize the enzymatic potential among species, CAZymes were identified from genomic sequences using run_dbCAN [49–51]. Protease detection was performed through homology-based searches of amino acid sequences against the MEROPS [52] database. As a result, comprehensive genome-wide catalogs of carbohydrate-active enzymes and peptidases were generated for each species. However, these annotation pipelines do not directly indicate whether a given enzyme has reported activity against ECM substrates.

**Table 1.** Genome metadata and annotation summary for the 11 *Bacteroides* species analyzed in this study.

| Species | Strain | Source | Accession | Genome length (bp) | Genes (n) | CAZymes (n, % of genes) | Proteases (n, % of genes) |
| --- | --- | --- | --- | --- | --- | --- | --- |
| *Bacteroides acidifaciens* | A40 | DSM 15896 | GCA_000613385.1 | 4,849,058 | 4,913 | 317 (6.45%) | 241 (4.9%) |
| *Bacteroides caccae* | VPI 3452A | ATCC 43185 | GCA_000169015.1 | 4,564,814 | 3,672 | 275 (7.49%) | 201 (5.5%) |
| *Bacteroides cellulosyticus* | CRE21 | DSM 14838 | GCA_025567135.1 | 3,614,006 | 2,948 | 214 (7.26%) | 169 (5.7%) |
| *Bacteroides fragilis* | NCTC 9343 | ATCC 25285 | GCA_001997325.1 | 5,210,800 | 4,330 | 268 (6.19%) | 196 (4.5%) |
| *Bacteroides intestinalis* | JCM 13265 | DSM 17393 | GCA_000172175.1 | 6,052,596 | 4,708 | 464 (9.86%) | 229 (4.9%) |
| *Bacteroides ovatus* | NCTC 11153 | ATCC 8483 | GCA_900107475.1 | 6,373,930 | 4,823 | 500 (10.37%) | 241 (4.9%) |
| *Bacteroides salyersiae* | WAL 10018 | ATCC BAA-997 | GCA_031172035.1 | 5,561,372 | 4,300 | 364 (8.47%) | 221 (5.1%) |
| *Bacteroides stercoris* | VPI B5-21 | ATCC 43183 | GCA_000154525.1 | 4,009,829 | 3,396 | 184 (5.42%) | 163 (4.8%) |
| *Bacteroides thetaiotaomicron* | VPI 5482 | ATCC 29148 | GCA_028539425.1 | 6,247,619 | 4,872 | 445 (9.13%) | 228 (4.7%) |
| *Bacteroides uniformis* | VPI 0061 | DSM 8492 | GCA_900107315.1 | 4,634,992 | 3,797 | 311 (8.19%) | 186 (4.9%) |
| *Bacteroides xylanisolvens* | XB1A | DSM 18836 | GCA_000210075.1 | 5,976,145 | 4,749 | 400 (8.42%) | 208 (4.4%) |

Thus, to enable systematic identification of ECM-targeting candidates, we constructed a substrate-driven ECM enzyme database. A list of 67 core human ECM terms [53, 54] spanning structural proteins, proteoglycans, glycoproteins, and glycosaminoglycans (**Supplementary Table 1**) was used to query the BRaunschweig ENzyme DAtabase (BRENDA) [55, 56]. All enzymes reported to interact with these substrates were retrieved and mapped to CAZyme family assignments and MEROPS identifiers. This curated database was then used to identify ECM-specific CAZYmes and proteases in each *Bacteroides* genome.

Across all genomes examined, both CAZymes and proteases predicted to interact with ECM substrates were detected (**Fig. 1**). ECM-targeting CAZyme genes (70 to 187 per genome) were more abundant than ECM-targeting protease genes (37 to 50 per genome) for all species analyzed (**Fig. 1A-B**). This predominance of CAZyme genes is consistent with the well-characterize role of *Bacteroides* species in glycan metabolism [23].

**Figure 1.**
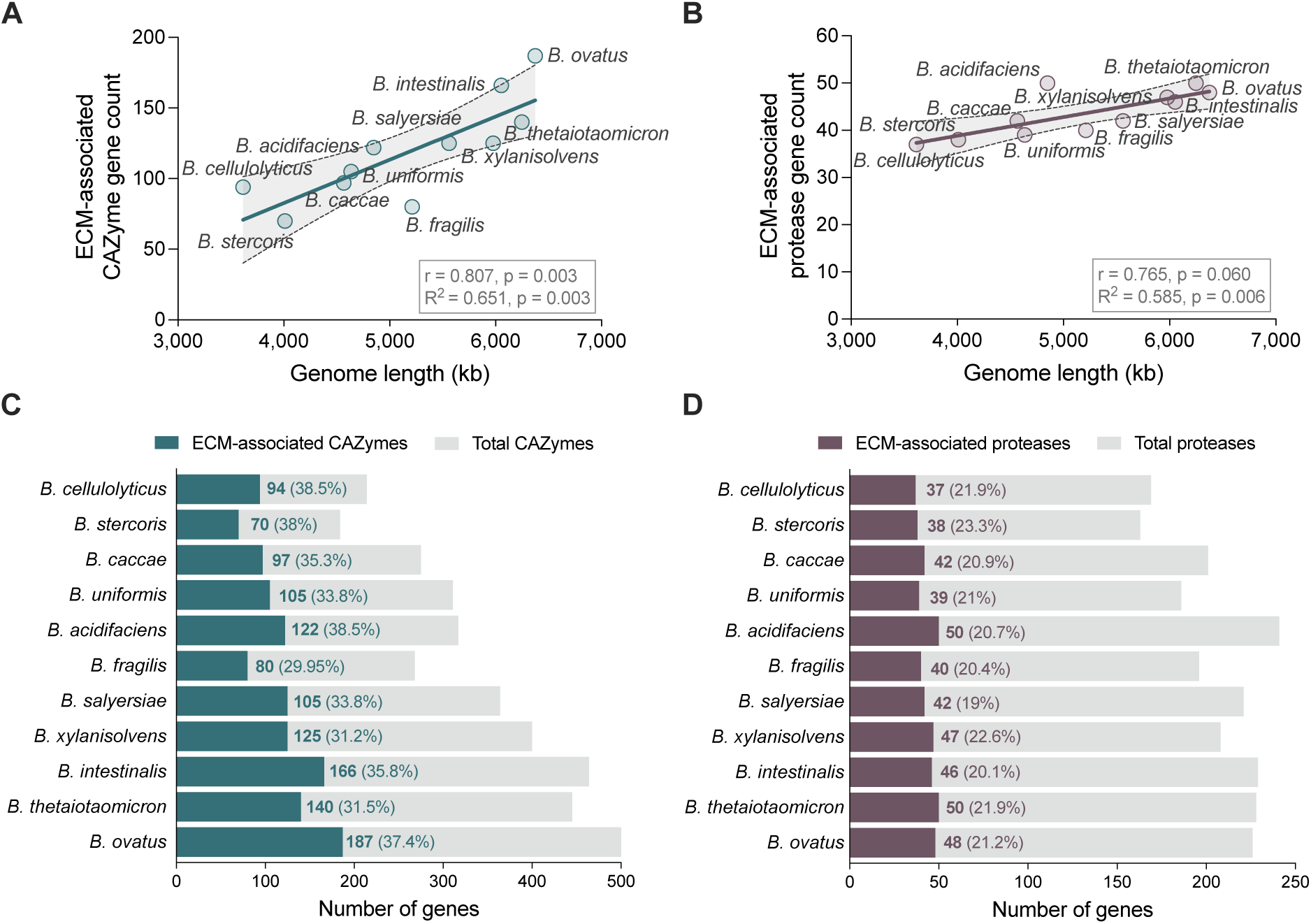
Abundance of ECM-associated CAZyme and protease genes across *Bacteroides* genomes. **(A-B)** Relationship between genome length and ECM-specific (A) CAZyme or (B) protease gene counts. Each point represents a species; the line indicates linear regression with 95% confidence intervals shown. The Pearson correlation coefficient (r), coefficient of determination (R^2^), and associated p value are displayed. **(C)** Total and ECM-specific CAZyme gene counts per species. **(D)** Total and ECM-specific protease gene counts per species. For (C-D), percentages indicate the proportion of total (C) CAZymes or (D) proteases predicted to target ECM substrates. Species are ordered by ascending genome length.

ECM-specific enzyme gene counts varied between species and were generally higher in genomes with greater total gene content (**Fig. 1A-B**, **Table 1**). ECM-specific CAZyme gene abundance exhibited a strong positive correlation with genome length (r = 0.807, p = 0.003; **Fig. 1A**), whereas ECM-specific protease gene abundance exhibited a weaker correlation with genome size (r = 0.765, p = 0.060; **Fig. 1B**). Similar positive correlations were observed between total CAZyme and total protease gene counts and genome size (**Supplementary Fig. 1**). When expressed relative to total enzyme content, ECM-specific CAZymes comprised 30-39% of total CAZymes (**Fig. 1C**) and ECM-specific proteases represented 19-23% of total proteases (**Fig. 1D**), indicating that ECM-targeting enzymes constitute a substantial fraction of both the CAZyme and protease repertoires across species. Together, these results demonstrate that enzymes with reported ECM-targeting activity are broadly encoded in *Bacteroides* genomes.

### *Bacteroides* genomes encode diverse and partially conserved ECM-associated CAZyme and protease families

Having established that ECM-targeting enzymes are broadly encoded across *Bacteroides* genomes, we assessed the functional diversity within this set of enzymes. For CAZymes, we first grouped these genes by CAZy class to capture broad catalytic categories. Five CAZy classes were represented: glycoside hydrolases (GH), carbohydrate-binding modules (CBM), polysaccharide lyases (PL), carbohydrate esterases (CE), and glycosyl transferases (GT; **Fig. 2A**). Across all species, GHs were the most abundant class, followed by CBMs and PLs, while CEs and GTs were present at lower levels. Because GH and PL families consist of catalytically active enzymes that cleave glycosidic linkages [49], their predominance indicates that ECM-associated CAZymes largely correspond to degradative functions. Protease genes were similarly grouped by catalytic type to capture broad mechanistic categories. ECM-associated proteases spanned six catalytic types, including serine, metallo-, cysteine, aspartic, asparagine, and threonine peptidases (**Fig. 2B**). Across all species, metallo- and serine-peptidases where the most abundant catalytic types, with cysteine proteases also consistently represented.

**Figure 2.**
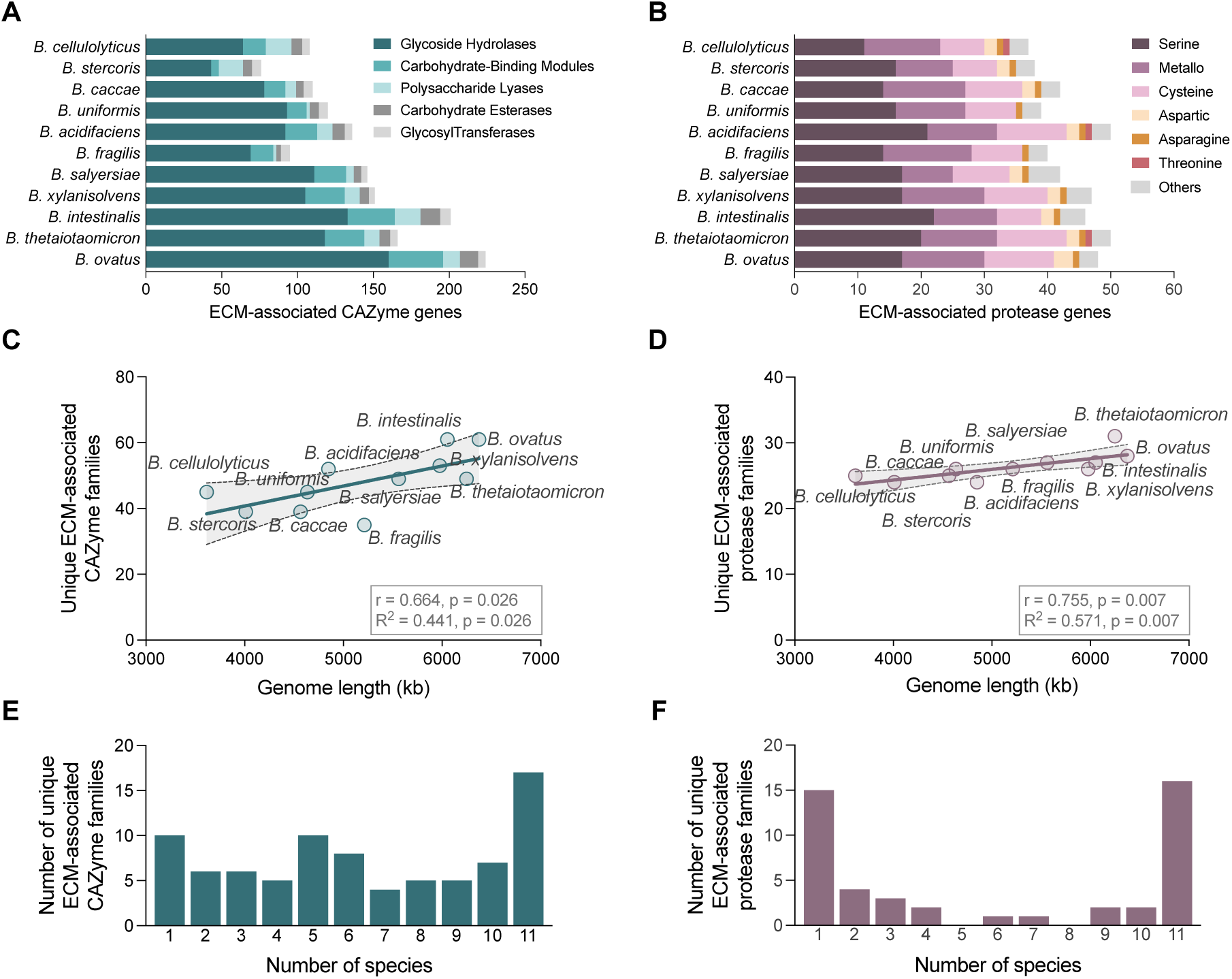
Distribution of ECM-associated CAZyme and protease classes and families. **(A)** Distribution of ECM-specific CAZyme genes grouped by CAZy class for each species. Because individual CAZymes may contain multiple functional domaines, genes can be represented in more than one class. **(B)** Distribution of ECM-specific protease genes grouped by catalytic type for each species. For (A-B), species are ordered by ascending genome length. **(C-D)** Relationship between genome length and the number of unique ECM-associated (C) CAZyme or (D) protease families per species. Each point represents a species; the line indicates linear regression with 95% confidence intervals shown. The Pearson correlation coefficient (r), coefficient of determination (R^2^), and associated p value are displayed. **(E-F)** Histogram displaying the prevalence of unique ECM-associate (E) CAZyme and (F) protease families across species. The x-axis indicated the number of species (1-11) in which a family is detected, and the y-axis indicates the number of families at each prevalence level.

To analyze functional diversity at a finer scale, we quantified the number and distribution of unique ECM-associated CAZyme and protease families across species. *Bacteroides* genomes encoded between 35-61 unique ECM-associated CAZyme families and 24-31 unique ECM-associated protease families. Family-level diversity varied across species and generally increased with genome length for both enzyme types (CAZymes: r =0.664, p = 0.026; proteases: r = 0.755, p=0.007; **Fig 2C-D**), although these correlations were more modest than those observed for gene-level abundance (**Fig. 1**). Furthermore, ECM-associated CAZymes exhibited a higher number of genes per family (2.44 ± 0.35 genes per family) than proteases (1.66 ± 0.16 genes per family), indicating greater within-family redundancy among CAZymes.

Next, we evaluated how these ECM-associated families were distributed across species (**Fig. E-F**). In total, 33 enzyme families (17 CAZyme, 16 protease) were detected in all 11 genomes, indicating a subset of predicted ECM-targeting families are shared across the genus. Beyond this conserved group, distinct distribution patterns emerged. ECM-associated CAZyme families were broadly represented across intermediate prevalence levels, with multiple families detected in 4-10 species (**Fig. 2E**). In contrast, protease families exhibited a more bimodal distribution, with 15 families restricted to a single species and fewer families represented at intermediate frequencies (**Fig. 2F**).

We also examined the gene copy number within individual CAZyme and protease families to further resolve these distribution patterns, (**Fig. 3**). Among CAZymes, a core set of 10 GH families was present in all *Bacteroides* species; these included GH2, GH3, GH13, GH20, GH29, GH33, GH77, GH92, GH95, and GH97 (**Fig. 3A**). In addition to these conserved GH families, the polysaccharide lyase family PL8 was also present in all genomes, and several CBM and GT families were broadly represented across species (**Supplementary Fig. 2**). Beyond this core subset, other GH, PL, CE, and GT families exhibited variable presence and gene copy number across species. A similar analysis was performed for protease families (**Fig. 3B**). A core set of 6 serine (S09, S13, S14, S16, S26, S54), 4 metallo- (M20, M24, M38, M49), 4 cysteine (C01, C26, C40, C44), and 1 asparagine (A02) were identified in all 11 genomes. The remaining proteases families exhibited substantial variation across genomes, with species-specific patterns of both presence and gene copy number.

**Figure 3.**
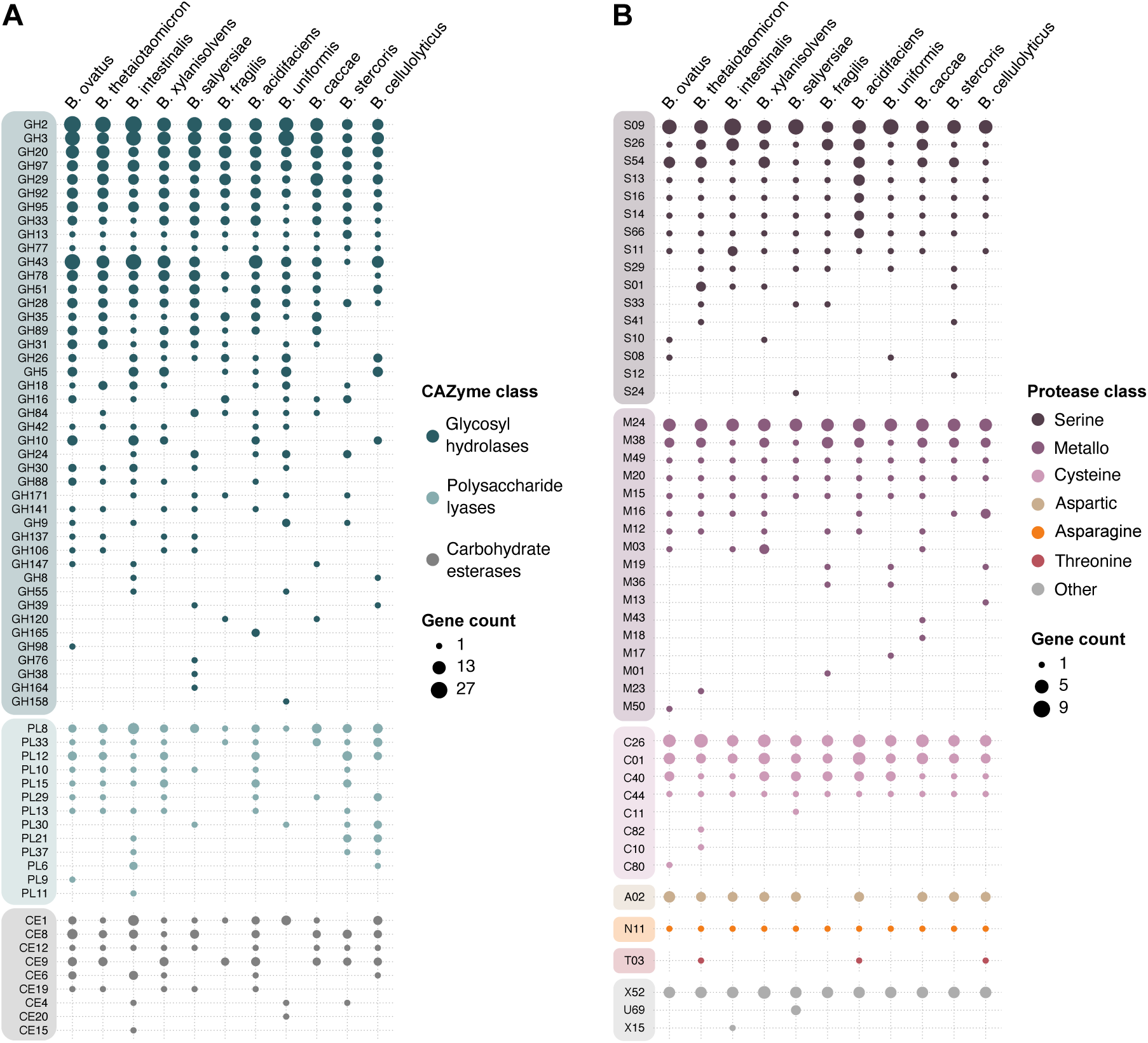
Family-level gene copy number of ECM-associated CAZymes and proteases. Dotplots of gene copy numbers for individual **(A)** CAZyme and **(B)** protease families. Rows represent individual CAZyme families grouped by (A) CAZy class or (B) catalytic type. Columns represent individual species. Dot size indicates gene copy number per family per species. Within each class, families are ordered first by decreasing prevalence across species and, among families with similar prevalence, by total gene copy number.

Collectively, these findings reveal that ECM-associated enzymatic capacity in *Bacteroides* is structured around a conserved set of CAZyme and protease families but is further shaped by species-specific variation in family composition and gene copy number. This diversification within and between genomes, particularly among CAZymes, suggests heterogeneous genomic potential for ECM modifications across species in the genus.

### Predicted substrate mapping reveals broad ECM coverage with enrichment for glycosaminoglycan-targeting enzymes and strain-level variation

In addition to the enzyme-centered analyses above, we quantified predicted ECM substrate coverage by mapping ECM-associated gene counts to their reported substrates and quantifying gene counts for individual components present in intestinal ECM (**Fig. 4; Supplementary Table 2**). Because both CAZymes and proteases can contribute to ECM modification, the substrate-level gene counts represent the integrated genomic capacity for degrading each ECM component. Across all species, the genomes encoded enzymes associated with numerous distinct ECM substrates, indicating broad predicted coverage rather than restriction to a single ECM class. Predicted targets spanned proteins, glycoproteins, proteoglycans, and glycosaminoglycans (GAGs).

**Figure 4.**
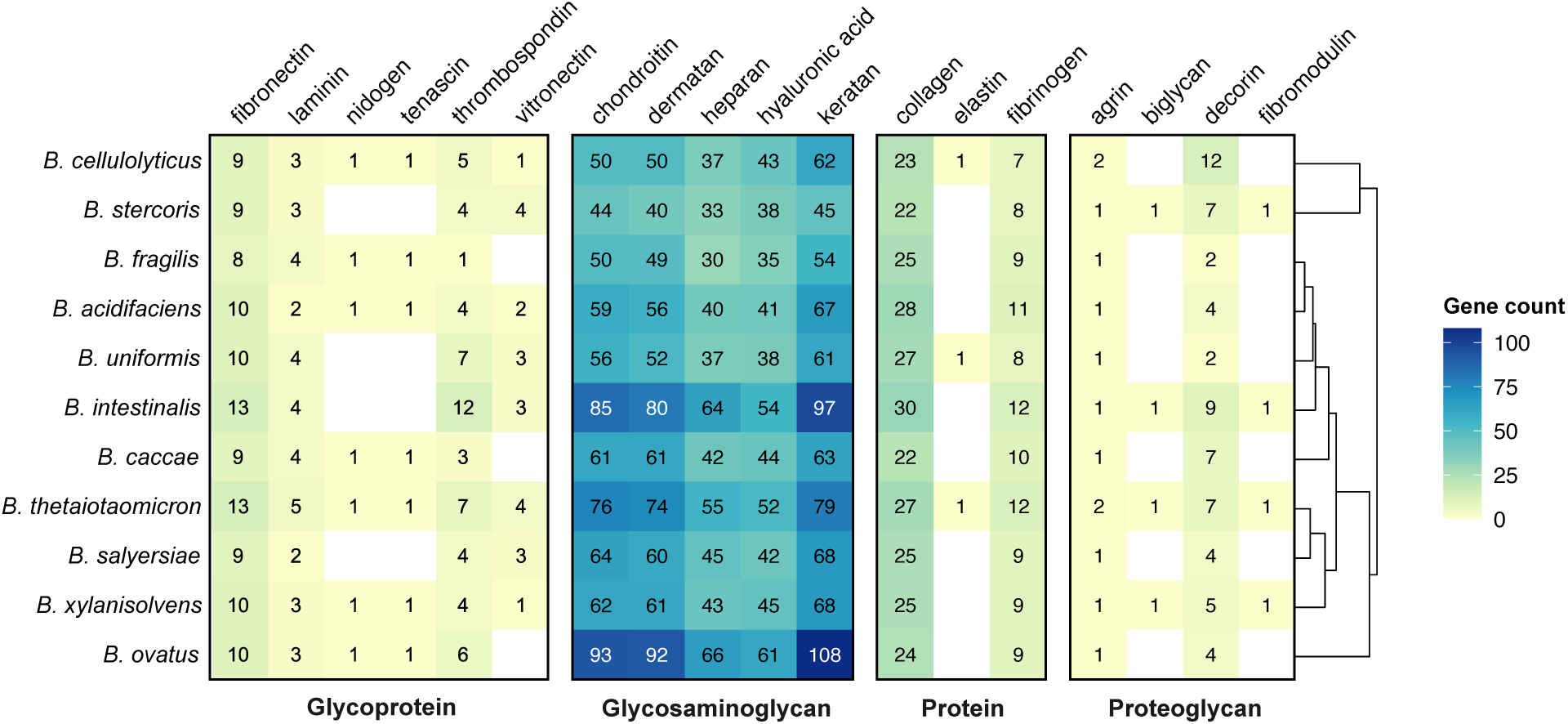
Substrate-level mapping of ECM-associated gene counts. Heatmap showing predicted ECM-associated gene counts mapped to individual extracellular matrix components. Rows represent species and columns represent ECM substrates grouped by category (glycoprotein, glycosaminoglycan, protein, and proteoglycan). Cell values indicate total gene counts per substrate per species; color intensity reflects gene count magnitude. Species are ordered according to hierarchical clustering based on Bray–Curtis dissimilarity of substrate-level relative gene count profiles (dendrogram shown at right).

For all species, GAG-associated genes comprised the largest proportion of gene counts, significantly exceeding those linked to protein substrates. Specifically, keratan (70 ± 18 genes per genome), chondroitin (63 ± 15), and dermatan (61 ± 15) sulfate were consistently associated with the highest gene counts across genomes, with genes targeting heparan sulfate (44 ± 12) and hyaluronic acid (45 ± 7) represented at comparatively lower levels. Among protein substrates, collagen-targeting genes (25 ± 2) were the most abundant, followed by fibrinogen (9 ± 2), the glycoprotein fibronectin (10 ± 2), and decorin (8 ± 3), a proteoglycan. These patterns are consistent with the well-established specialization of Bacteroides species in glycan foraging [57, 58] and suggest that intestinal ECM glycosaminoglycans represent a dominant predicted substrate class within this genus.

To visualize species-level similarity in substrate composition, we clustered genomes using Bray–Curtis dissimilarity calculated from relative substrate-level gene abundances (**Fig. 4**; **Supplementary Fig. 3**). The resulting dendrogram revealed organized species into several clusters, including a close association between *B. cellulolyticus* and *B. stercoris* and between *B. salyersiae* and *B. xylanisolvens*. Overall, however, species did not segregate into distinct substrate-specialized groups, consistent with broadly shared ECM substrate coverage across the genus.

### Experimental assessment of ECM degradation reveals substrate- and species-specific activity patterns

To determine whether the predicted ECM-associated gene repertoires translate into measurable enzymatic activity, we quantified the degradation of several intestinal ECM substrates using supernatants collected from cultures of the same 11 *Bacteroides* species. We evaluated degradation of both glycosaminoglycans (chondroitin sulfate, heparan sulfate, and hyaluronic acid) and proteins (gelatin – denatured collagen, collagen IV, fibronectin, and elastin). Across the substrates tested, most species exhibited detectable degradation in at least one assay, and each substrate was degraded by at least four species, indicating that ECM-degrading activity is broadly distributed across the *Bacteroides* genus, consistent with the widespread genomic encoding of ECM-associated CAZymes and proteases described above (**Fig. 5**).

**Figure 5.**
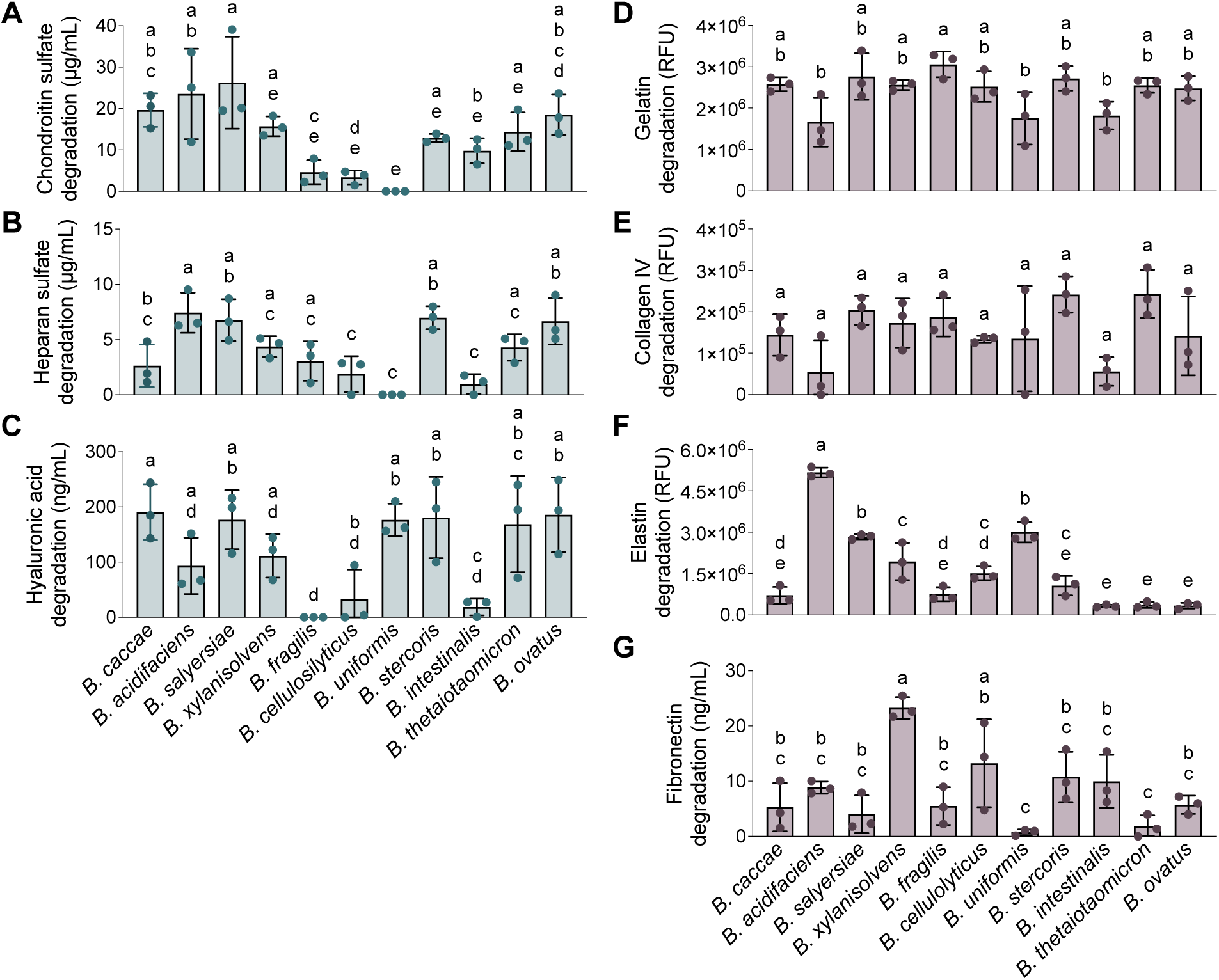
Experimental assessment of ECM substrate degradation by *Bacteroides* supernatants. Degradation of extracellular matrix substrates by culture supernatants from 11 *Bacteroides* species. (**A–C**) Glycosaminoglycan substrates: chondroitin sulfate, heparan sulfate, and hyaluronic acid. (**D–G**) Protein substrates: gelatin (denatured collagen), collagen IV, elastin, and fibronectin. Values were corrected by subtracting media-only controls. Bars represent mean ± standard deviation of three biological replicates, each consisting of the average of three technical replicates. Statistical significance was determined by one-way ANOVA followed by Tukey’s multiple comparison test; different letters indicate significant differences among species (p < 0.05). Species are ordered in the x-axis according to the hierarchical clustering in Figure 4.

GAG degradation was widespread but uneven across species (**Fig. 5A-C**). Several species, including *B. stercoris, B. thetaiotaomicron, B. salyersiae, B. xylanisolvens*, and *B. ovatus,* exhibited relatively high degradation across all three GAG substrates. In contrast, *B. cellulolyticus* and *B. intestinalis* demonstrated limited activity against these substrates. Other species displayed substrate-specific patterns. For example, *B. fragilis* did not degrade hyaluronic acid despite showing measurable activity toward chondroitin and heparan sulfate, and *B. uniformis* displayed high activity against hyaluronic acid but no degradation of the other two GAG substrates.

Degradation of protein ECM substrates also varied across species with distinct substrate-dependent patterns (**Fig. 5D-G**). Gelatin and collagen IV were degraded by most species at similar magnitudes, indicating that collagen-targeting proteolytic activity is broadly distributed across the genus (**Fig. 5D-E**). In contrast, elastin degradation was restricted to a subset of species, with significant activity observed in *B. cellulolyticus, B. acidifaciens, B. uniformis, B. salyersiae,* and *B. xylanisolvens,* and the rest of the species displaying minimal enzymatic activity (**Fig. 5F**). Fibronectin degradation was also heterogenous across species, with *B. xylanisolvens* and *B. cellulolyticus* among the high degraders and several species exhibiting low activity (**Fig. 5G**).

Together, these findings indicate that while many *Bacteroides* species possess the capacity to degrade multiple ECM components, the extent and substrate specificity of this activity differ across species, reflecting functional diversification within the genus.

### Genomic encoding and enzymatic activity exhibit limited concordance

Given the broad yet species-specific degradation patterns observed experimentally, we examined the extent to which substrate-level gene abundance aligned with measured enzymatic activity. Spearman correlation analyses between substrate-level gene counts and degradation magnitude revealed no significant correlations for any ECM substrate tested (**Supplementary Fig. 4A**). We next assessed whether overall species similarity patterns were conserved between datasets by comparing genomic and experimental distance matrices using a Mantel test. This analysis revealed no significant concordance (Mantel r = -0.152, p = 0.764) between species-level genomic profiles and their corresponding enzymatic activity patterns.

Principal component analysis (PCA) was used to further compare the multivariate structure of the substrate-level genomic and experimental datasets (**Fig. 6**). In the genomic PCA (**Fig. 6A**), separation along PC1 (62.6% of the variance) was primarily driven by overall gene abundance across genomes, whereas PC2 (21%) reflected differences in the relative contribution of GAG-versus protein-associated gene counts (**Supplementary Fig. 4B**). In the experimental PCA (**Fig. 6B**), species segregated according to distinct degradation activity patterns across substrates. An analysis of substrate loadings revealed that PC1 (35.6% of the variance) was driven primarily by GAG degradation activity, while PC2 (25%) reflected differences associated with collagen and elastin degradation activity (**Fig. 6C**). Thus, although GAG-associated features contributed prominently to variation in both datasets, species segregation in the experimental PCA did not mirror that observed in the genomic PCA.

**Figure 6.**
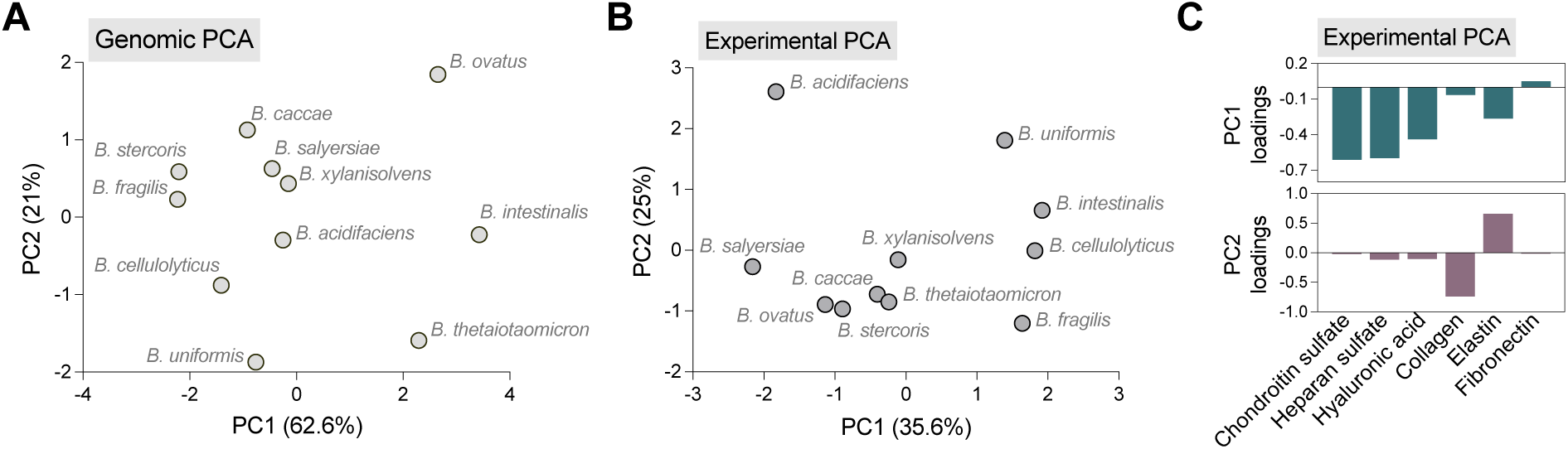
Principal component analysis of substrate-level gene counts and experimental ECM degradation activity. (**A**) PCA of substrate-level ECM-associated gene counts across 11 *Bacteroides* species using only the ECM substrates evaluated experimentally. Gene counts were Z-score normalized prior to analysis. (**B**) PCA of ECM degradation activity measured in bacterial culture supernatants. Degradation values were Z-score normalized prior to analysis. (**C**) Substrate loadings for the experimental PCA shown in (B). Bars indicate the contribution of each ECM substrate to PC1 and PC2.

### Protease inhibition confirms contributions of multiple catalytic types to collagen degradation

Finally, we evaluated whether the protease catalytic classes identified in the genomic analysis (**Fig 2B**) were functionally involved in ECM degradation. Collagen was selected as a representative ECM protein substrate because of its prominence in intestinal tissue, its relatively high gene abundance compared to other protein substrates, and its robust degradation across most species in our experimental assays. First, we examined the catalytic type distribution of predicted collagen-targeting proteases across *Bacteroides* genomes (**Fig. 7A**). Collagen-associated proteases spanned multiple catalytic classes, including serine, metallo-, and cysteine peptidases, with serine and metalloproteases representing the most abundant predicted classes in all species.

**Figure 7.**
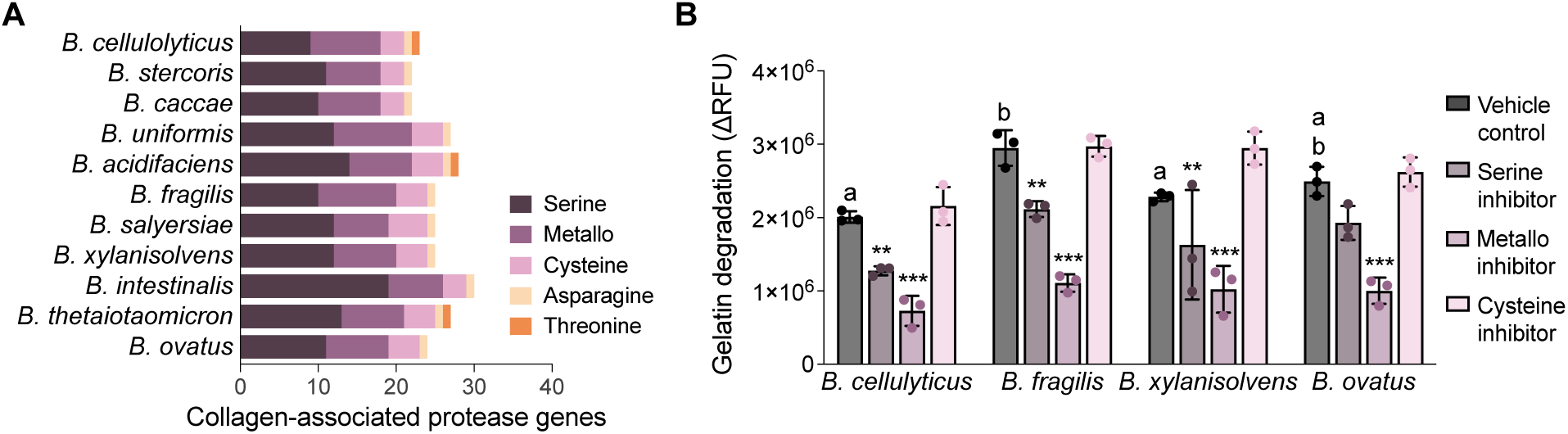
Functional assessment of protease classes contributing to gelatin degradation. **(A)** Distribution of predicted collagen-associated protease genes grouped by catalytic type across *Bacteroides* genomes. Species are ordered by ascending genome length. **(B)** Quantification of gelatin degradation in the presence of inhibitors targeting serine, metallo-, and cysteine proteases in four representative species. Bars represent mean ± standard deviation of three biological replicates, each consisting of the average of three technical replicates. Statistical significance was determined by two-way ANOVA followed by Tukey’s multiple comparisons test. Vehicle controls were compared across species – different letters denote statistical significance (p<0.05). Asterisks indicate significant differences relative to the vehicle control for that species (*p<0.05, **p<0.01, ***p<0.001).

We then tested these predicted catalytic contributions by quantifying gelatin degradation in the presence of inhibitors targeting serine, metallo-, and cysteine proteases (**Fig. 7C**.) Metalloprotease inhibition produced the most consistent and pronounced reduction in gelatin degradation across the four species tested, resulting in a 60-74% decrease in activity. Serine protease inhibition also led to statistically significant reductions (23-39%) in all supernatants except *B. ovatus.* In contrast, cysteine protease inhibition did not significantly alter gelatin degradation under the conditions tested, despite the presence of predicted cysteine collagen-targeting proteases in these genomes. These results highlight that genomic presence alone does not necessarily predict measurable catalytic activity. Furthermore, no single inhibitor completely abolished gelatin degradation, confirming that multiple protease classes likely contribute to collagenolytic activity.

## DISCUSSION

The *Bacteroides* genus is well established as a dominant and metabolically versatile component of the human gut microbiome, renowned for its extensive repertoire of CAZymes and capacity to degrade structurally complex host- and diet-derived glycans [59–61]. Here, we extend this metabolic understanding by demonstrating that ECM-targeting enzymatic activity, spanning both proteolytic and glycan-degrading functions, is a broadly encoded and functionally active trait across the genus. Through comparative genomic profiling of 11 species combined with substrate-specific degradation assays, we show that *Bacteroides* encodes diverse CAZymes and proteases capable of targeting the major components of the intestinal ECM, including collagen, elastin, fibronectin, and GAGs. While our prior work identified several *Bacteroides* species as particularly active ECM degraders among a broader panel of gut commensals [31], the present study reveals that this capacity is conserved across the genus.

The widespread encoding of ECM-targeting enzymes across all 11 *Bacteroides* species examined here indicates that this capacity is a conserved genus-level feature rather than an isolated phenotype restricted to a few species. Nevertheless, the repertoire of ECM-associated CAZymes and proteases varied considerably across genomes, with species such as *B. ovatus, B. thetaiotaomicron,* and *B. intestinalis* encoding substantially broader enzymatic arsenals than *B. stercoris* or *B. fragilis*. Inter-species heterogeneity correlated with overall genome size, consistent with the well-documented relationship between genome expansion and metabolic versatility in this genus [33, 34, 61]. This correlation was stronger for ECM-associated CAZymes than proteases, reflecting the crucial role of polysaccharide foraging as a driver of genome diversification in *Bacteroides* [23, 62]. Indeed, *Bacteroides* genomes are highly plastic, shaped by lateral gene transfer, gene duplication, and mobile genetic elements, which have been linked to the acquisition of new carbohydrate-degrading capabilities and niche adaption [63–65].

The abundance of ECM-associated CAZymes across *Bacteroides* genomes was largely attributable to enzymes predicted to target GAG substrates, which primarily included glycoside hydrolases and polysaccharide lyases. This was mirrored experimentally by broad degradation activity in the genus against chondroitin sulfate, heparan sulfate and hyaluronic acid. Foundational biochemical and genomic characterization of GAG metabolism in *B. thetaiotaomicron* has established that degradation of these substrates is mediated by dedicated polysaccharide utilization loci (PULs) encoding polysaccharide lyases, glycoside hydrolases, and sulfatases [66, 67]. Building on this, Overbeeke *et al.* demonstrated that the capacity to degrade chondroitin sulfate and hyaluronic acid varies across *Bacteroides* species and is shaped by PUL architecture [68]. Species-specific degradation patterns in our experimental data aligned with prior independent characterizations: high chondroitin sulfate activity by *B. salyersiae* [48] and *B. thetaiotaomicron* [42, 43, 67], heparan sulfate degradation by *B. stercoris* [69], and variable hyaluronic acid activity across species, with generally low activity by *B. fragilis* and *B. intestinalis* [41, 47].

Turning to protein substrates, proteolytic ECM degradation in the *Bacteroides* genus is best characterized in enterotoxigenic *B. fragilis* strains, whose pathogenicity island encodes fragilysin, a zinc metalloprotease with documented gelatinase and type IV collagenase activity [39, 40]. Beyond fragilysin, biochemical characterization of *B. fragilis* supernatants has identified serine, metallo- and cysteine proteases active against gelatin and fibrinogen [70–72]. More recently, we demonstrated that *B. fragilis, B. thetaiotaomicron,* and *B. ovatus,* among a panel of gut commensals, degraded multiple ECM substrates, including GAGs, collagen, laminin, and fibronectin [31]. The present study builds on these findings by demonstrating that collagenolytic activity is broadly distributed across the genus, consistent with earlier surveys identifying gelatinase and collagenase activities in clinical *Bacteroides* isolates [73]. Inhibitor experiments confirmed that collagenolytic activity is driven primarily by metalloproteases and secondarily by serine proteases across representative species. Fibronectin degradation, previously reported for *B. fragilis* [31, 74] and *B. ovatus* [31] was also detected across multiple species here, while elastin degradation, largely undocumented in intestinal *Bacteroides* [75], emerged as a capacity restricted to a subset of species, representing a novel observation for the genus.

The limited concordance between genomic predictions and experimental ECM degradation activity observed here is consistent with the well-recognized gap between genotype and phenotype in microbial systems [76]. Several factors likely contribute to this divergence. From a computational standpoint, our approach relies on substrate-enzyme associations curated in databases whose experimentally characterized entries are predominantly derived from eukaryotic organisms [77]. Furthermore, EC number classification captures reaction type rather than enzyme identity, meaning enzymes sharing the same EC number can exhibit substantially different substrate specificities [78]. These limitations likely also result in false negatives, as novel ECM-degrading enzymes lacking characterized homologs would escape detection entirely. Experimentally, our assays were conducted under conditions that may not fully recapitulate *in vivo* enzymatic activity. For example, cysteine and aspartic proteases exhibit optimal activity at acidic pH, whereas serine proteases and metalloproteases are active at neutral pH [79], potentially explaining why cysteine proteases inhibition had no measurable effect on gelatin degradation in our neutral pH assays despite genomic predictions. Finally, our assays employed soluble ECM substrates, whereas native intestinal ECM exists in crosslinked fibrillar form; enzymatic activity against matrix-form substrates may differ substantially from the solution-phase assays employed here [80].

*Bacteroides* species have been associated with intestinal diseases involving aberrant ECM remodeling [36–38], raising the question of whether the enzymatic capacity characterized here contributes to disease pathogenesis. A first consideration is how commensal bacteria usually present in the lumen gain access to host ECM. *Bacteroides* constitutively produce outer membrane vesicles that can deliver enzymatic cargo to host cells independently of direct bacterial contact [42, 81]. Additionally, increased epithelial permeability in disease facilitates both bacterial translocation and diffusion of secreted enzymes into the subepithelial space [82–86]. Once in proximity to ECM, microbial enzymatic activity could amplify inflammation by generating bioactive matrix fragments that may modulate immune cell behavior [87–90]. Consistent with this possibility, proteolytic activity attributed to members of the order Bacteroidales has been linked to ulcerative colitis onset and severity [24, 91, 92], inflammatory bowel disease progression [93, 94], and anastomotic leakage following colorectal surgery [95].

Collectively, our findings establish *Bacteroides* as a genus with broad, functionally active ECM-degrading capacity spanning both proteolytic and glycan-degrading functions, positioning commensal gut bacteria as previously underappreciated contributors to intestinal ECM remodeling. Future work examining clinical isolates and disease-associated strains will be essential to determine how this capacity varies beyond type strains and whether it scales with disease severity. Ultimately, incorporating microbial enzymatic activity into the conceptual framework of intestinal ECM homeostasis opens new avenues for understanding how the gut microbiome shapes tissue architecture and for designing microbiota-targeted strategies to modulate remodeling in the context of intestinal disease.

## METHODS AND MATERIALS

### Curation of the ECM enzyme database

We generated a comprehensive ECM enzyme database using a substrate-based approach. A list of 67 core human ECM terms [53, 54] was used to query and import the BRENDA-enzyme database (accessed on 3/28/2025). The list of ECM components represents various broad categories including proteins, proteoglycans, and glycoproteins (**Supplementary Table 1**). All enzyme names and EC numbers reported to interact with each substrate were retrieved from BRENDA using the R package brendaDb [56]. Each retrieved EC number was mapped to enzyme classification databases using the UniProtKB API call [96, 97], yielding CAZyme family and subfamily assignments for carbohydrate-active enzymes and MEROPS identifiers for peptidases. The resulting curated database (full dataset available at https://github.com/PorrasTMILab/CMBE-Bacteroides-ECM) contains enzyme names, EC numbers, CAZyme family assignments, and MEROPS identifiers for all enzymes reported to interact with each ECM substrate based on the primary literature entries that inform the BRENDA database.

### Functional genome annotation and identification of ECM-specific enzymes

Type strain reference genomes from 11 *Bacteroides* species were obtained from the National Center for Biotechnology Information RefSeq database (accessed on 05/05/2025). The analyzed species include: *B. acidifaciens*, *B. caccae*, *B. cellulolyticus*, *B. fragilis*, *B. intestinalis*, *B. ovatus*, *B. salyersiae*, *B. stercoris*, *B. thetaiotaomicron*, *B. uniformis*, and *B. xylanisolvens* (source and strain information available in **Table 1**). We performed structural annotation of the assembled sequences using Prokka (version 1.14.6) [98]. Then, the predicted gene sequences from Prokka were functionally annotated using eggNOG-mapper (version emapper-2.1.12) [99–101], which utilizes DIAMOND [101] sequence homology searches (e-value threshold of 0.001) to assign gene functions and enzyme commission (EC) numbers.

For CAZyme identification, fully assembled genome sequences were processed using run_dbCAN (version 5.0.0) [49–51] with default prokaryote analysis parameters. Within run_dbCAN, gene prediction was performed by Prodigal (version 5.1.2) and annotations were assigned using DIAMOND BLAST [101] against the Carbohydrate Active Enzymes database (http://www.cazy.org/) [102]. Additionally, pyHMMER [103, 104] was used to search against the dbCAN-HMM and dbCAN_sub-HMM profiles for enhanced protein domain detection. Only CAZyme annotations supported by at least two independent methods were retained for downstream analysis.

To identify proteases, we conducted a homology search using the amino acid sequences predicted by Prokka against the MEROPS Peptidase Database [52] (release version 12.4, downloaded https://www.ebi.ac.uk/merops/ on 05/01/2025) using phmmer (HMMER version 3.4) with an e-value threshold of 0.01. Genes that matched to an inhibitor or a non-peptidase homologue were excluded from further analysis.

Next, the curated ECM enzyme database was used to identify ECM-specific CAZymes and proteases within all annotated *Bacteroides* CAZymes and proteases by matching EC numbers and MEROPS family assignments. Where available, EC numbers from run_dbCAN annotations were used; otherwise, EC numbers from eggNOG annotations were substituted. For inclusion in the ECM-specific gene counts, genes had to be confirmed as CAZymes by run_dbCAN or as proteases by the MEROPS search. Gene counts were calculated as the total number of unique gene IDs per species, substrate, family, or class.

### Bacterial culture and supernatant collection

Reference strains were obtained from the American Type Culture Collection and Leibniz Institute DSMZ (**Table 1**). All bacteria were cultured in brain heart infusion broth (BHIS) supplemented with 5 µg/mL hemin and 1 mg/mL L-cysteine. For agar plates, an additional 15 g/L agar powder was added to the solution. Cultures were maintained in a vinyl anaerobic chamber (Coy Lab Products) at 37°C. Frozen glycerol stocks of each species were streaked on agar plates, single colonies were picked after 48 hours and inoculated in liquid cultures. Each colony grew overnight in broth until reaching stationary phase, then fresh BHIS was inoculated with the overnight culture at a 1:100 dilution. Once the cultures reached an OD600 of 0.65-0.75, they were centrifugated at 6,000 x g for 10 minutes at 4°C to isolate supernatants. After supernatant collection, aliquots were stored at -20°C until use. Routine verification of species identity was performed using Sanger sequencing for the V4 region of the 16S rRNA gene using 515F/806R primers [105]. Supernatants were collected from three individual runs with three technical replicates per run. Routine verification of species identity was performed using Sanger sequencing for the V4 region of the 16S rRNA gene using 515F/806R primers [105]. Supernatants were collected from three individual runs with three technical replicates per run.

### *In vitro* assessment of ECM proteins degradation

The total protein concentration in the bacterial culture supernatants was measured using a Pierce Micro BCA assay kit (23235, Thermo Fisher) following the manufacturer’s protocol. Supernatants were diluted in cold PBS to a final concentration of 10 mg/mL. A broth control was also prepared consisting of BHIS cultured in the anaerobic chamber for the same amount of time as the supernatants. Substrate-based assays were performed to measure degradation of individual ECM components by the bacterial culture supernatants.

Elastin degradation was quantified using a fluorometric commercial kit (EnzChek Elastase Assay Kit, E12056, Invitrogen) according to the manufacturer’s instructions with minor modifications. Briefly, 10 mg/mL protein concentration of bacterial culture supernatant was incubated with 25 ug/mL of DQ elastin substrate working solution diluted in 1x reaction buffer. After 4 hours of incubation at 37°C, the fluorescence intensity of the solutions was measured in a microplate plate reader (SpectraMax iD3, Molecular Devices) at an excitation/emission wavelength of 485 nm/535 nm. Each biological replicate represents the average of three technical replicates. Fluorescence values were normalized by subtracting the BHIS baseline controls.

Degradation of gelatin (D12054, Invitrogen), and collagen IV (D12052, Invitrogen) substrates were measured using fluorometric commercial kits, following the manufacturer’s protocol with some modifications. First, 1 mg/mL stock DQ gelatin or DQ collagen IV were diluted in the bacterial supernatants at a protein concentration of 10 mg/mL for a final concentration of 5% v/v DQ substrate solution. Then, the solutions were incubated for 8 hours at 37°C. Fluorescence was measured at the beginning and at the endpoint using an excitation/emission wavelength of 485 nm/525 nm in a plate reader (SpectraMax iD3, Molecular Devices). Fluorescence values of three biological replicates, each as the average of three technical replicates, were normalized by subtracting the baseline measurement at time zero from the corresponding measurement after 8 hours.

Fibronectin degradation was quantified using a human fibronectin ELISA kit (ELH-FN1, RayBiotech following the kit’s protocol. Bacterial supernatants at a protein concentration of 10 mg/mL were incubated for 24 hours at 37°C with 2 µg/L human fibronectin (FC010, Sigma-Aldrich) in phosphate buffer saline for a final concentration of 100 ng/mL fibronectin in supernatant. Absorbance was measured (λ = 450 nm) using a SpectraMax plate reader. The amount of intact fibronectin after incubation was inferred using a standard curve of human fibronectin protein. Degraded fibronectin values were calculated by subtracting the final fibronectin concentrations in supernatants from the final concentrations found in BHIS control.

### *In vitro* Characterization of Glycosaminoglycan Degradation

To assess hyaluronic acid (HA) degradation, bacterial supernatants at 10 mg/mL protein concentration were incubated with HA sodium salt (AAJ6699303, Fisher Scientific) at a final concentration of 200 ng/mL over 24 hours at 37°C. The final concentration of HA was measured using a Hyaluronan DuoSet ELISA (DY3614, R&D Systems) according to the manufacturer’s instructions. Readings was measured at a wavelength of 450 nm and the concentrations of HA were determined through the kit’s HA standard curve. Amounts of degraded HA are depicted as the difference between the BHIS control and the concentrations in each supernatant after incubation.

The degradation of sulfated GAGs was quantified using the 1,9-dimethyl-methylene blue (DMMB) assay [106]. The degradation of sulfated GAGs was quantified using the 1,9-dimethyl-methylene blue (DMMB) assay [106]. Chondroitin sulfate (CS; AAJ6034106, Fisher Scientific) and heparan sulfate (HS; HY-101916, MedChemExpress) stocks of 10 mg/mL were prepared in deionized water. Stock solutions of CS and HS were added to the diluted bacterial supernatants at a final concentration of 40 and 20 µg/mL, respectively, and incubated at 37°C for 8 hours. Standards are prepared in BHIS broth with concentrations ranging from 100 µg/mL to 0 µg/mL. After substrate incubation, 20 uL of each sample or standard were mixed with 200 uL DMMB dye reagent (16 mg/L in diH_2_O with 2.37 g sodium chloride and 3.04 g glycine; Sigma) in a clear 96-well plate. Absorbance was measured immediately on a plate reader (SpectraMax iD3, Molecular Devices) at a wavelength of 525 nm. Concentrations of the substrate were interpolated from the standard curve. Final degradation values were normalized by subtracting concentrations in the supernatant from the measured concentration in the media control.

### Inhibition of gelatin degradation

Proteolytic activity in DQ gelatin was inhibited using the same experimental conditions described previously, with the addition of a preincubation step. First, bacterial supernatants were preincubated for 20 minutes at room temperature with a different inhibitor dissolved in DMSO prior to the addition of each substrate. Enzymatic activity was inhibited for cysteine peptidases using 100 nM E-64 (HY-15282, MedChemExpress), serine peptidases with 200 µM AEBSF hydrochloride (HY-12821, MedChemExpress), and metallopeptidases with 1 mM 1,10-Phenantrholine monohydrate (HY-Y1841, MedChemExpress). The vehicle control was prepared by omitting the inhibitors and using an equivalent volume of DMSO.

### Comparative multivariate analysis of genomic and experimental activity

For each substrate, Spearman’s rank correlations were calculated between the genomic analysis (gene counts) and the experimental activity (mean activity) using the *Hmisc* [107] R package. Per-substrate correlation coefficients and p-values were computed using non-parametric methods. We applied Benjamini–Hochberg false discovery rate correction to adjust for multiple hypothesis testing.

A Mantel test was also performed to assess the correlation between the genomic prediction matrix (containing gene counts per substrate for each species) and the experimental observations matrix (comprising mean enzyme activities per substrate and species). Prior to analysis, both matrices were standardized by z-score normalization. Euclidean distance matrices were calculated from the normalized data to serve as inputs to the Mantel test. The Mantel test used Spearman’s rank correlation with permutation-based significance (9,999 permutations). The *vegan* [108] R package was used for the Mantel test and Euclidean distance calculations.

Principal component analysis (PCA) was conducted on the z-scored genomic predictions and experimental data matrices in Rstudio (version 2026.1.0.392) [109]. PCA scores and loadings were obtained using the *prcomp* R function and exported for visualization in in GraphPad Prism version 10.6.0 (GraphPad Software).

### Statistical analysis

Linear correlations between genome size and total gene counts, protease or CAZyme gene count, ECM-specific enzyme counts, or ECM-specific protease and CAZyme families were assessed using the Pearson correlation coefficient in GraphPad Prism version 10.6.0 (GraphPad Software). For the *in vitro* degradation and inhibition experiments, statistical significance was determined in GraphPad using one-way or two-way ANOVA, respectively, followed by Tukey’s multiple comparison test, assuming a normal Gaussian distribution. A threshold of P ≤ 0.05 was used to determine significance.

To explore and visualize the patterns in similarity among the ECM-specific gene profiles across species, we performed hierarchical clustering based on relative abundances of substrate-specific gene counts. Hierarchical clustering with average linkage was determined through pairwise Bray-Curtis dissimilarities from relative abundance matrices of ECM-specific gene counts. First, a matrix was constructed with gene counts for each species per substrate relevant to the intestine (**Supplementary Table 2**). The matrix was converted to relative abundance values by dividing each substrate gene count by the sum of all substrate gene counts for that species. Then, the Bray-Curtis dissimilarities were calculated using the vegdist function of the *vegan* R package [108].

## SUPPLEMENTARY INFORMATION

Supplementary figures and tables are provided in the supplementary file.

## DATA AND CODE AVAILABILITY

Data processing, analyses, and some figure generation were performed in R. Code scripts, and associated datasets are available through GitHub (https://github.com/PorrasTMILab/CMBE-Bacteroides-ECM).

## AUTHOR CONTRIBUTIONS

Conceptualization and methodology: KMMA and A.M.P; data curation: KMMA, BD, and IY; investigation: KMMA, KR, and MJ; formal analysis, visualization, and writing: KMMA and AMP; supervision and funding acquisition: A.M.P. All authors revised and approved the manuscript.

## ETHICS DECLARATIONS

### Ethical approval

Not applicable.

### Consent to participate

Not applicable.

### Consent to publish

Not applicable.

## REFERENCES

1. McKee, T. J., G. Perlman, M. Morris, and S. V. Komarova. Extracellular matrix composition of connective tissues: a systematic review and meta-analysis. Sci Rep Nature Publishing Group; 9(1):10542 2019. 10.1038/s41598-019-46896-0

2. Urbanczyk, M., S. L. Layland, and K. Schenke-Layland. The role of extracellular matrix in biomechanics and its impact on bioengineering of cells and 3D tissues. Matrix Biology 85–86:1–14 2020. 10.1016/j.matbio.2019.11.005

3. Lee, S.-E., I. Massie, L. Meran, and V. S. W. Li. Extracellular Matrix Remodeling in Intestinal Homeostasis and Disease. Advances in Stem Cells and their Niches Elsevier; 2:99–140 2018. 10.1016/BS.ASN.2018.01.001

4. Pompili, S., G. Latella, E. Gaudio, R. Sferra, and A. Vetuschi. The Charming World of the Extracellular Matrix: A Dynamic and Protective Network of the Intestinal Wall. Frontiers in Medicine Frontiers Media S.A.; 8:477 2021. 10.3389/FMED.2021.610189/BIBTEX

5. Jyoti, and P. Dey. Mechanisms and implications of the gut microbial modulation of intestinal metabolic processes. npj Metab Health Dis Nature Publishing Group; 3(1):24 2025. 10.1038/s44324-025-00066-1

6. Vilardi, A., S. Przyborski, C. Mobbs, A. Rufini, and C. Tufarelli. Current understanding of the interplay between extracellular matrix remodelling and gut permeability in health and disease. Cell Death Discov. Nature Publishing Group; 10(1):258 2024. 10.1038/s41420-024-02015-1

7. Lu, P., K. Takai, V. M. Weaver, and Z. Werb. Extracellular Matrix Degradation and Remodeling in Development and Disease. Cold Spring Harb Perspect Biol Cold Spring Harbor Lab; 3(12):a005058 2011. 10.1101/cshperspect.a005058

8. DeLeon-Pennell, K. Y., T. H. Barker, and M. L. Lindsey. Fibroblasts: The arbiters of extracellular matrix remodeling. Matrix Biology 91–92:1–7 2020. 10.1016/j.matbio.2020.05.006

9. Vizovišek, M., M. Fonović, and B. Turk. Cysteine cathepsins in extracellular matrix remodeling: Extracellular matrix degradation and beyond. Matrix Biology 75–76:141–159 2019. 10.1016/j.matbio.2018.01.024

10. Zhu, Y. et al. Interplay between Extracellular Matrix and Neutrophils in Diseases. Journal of Immunology Research Hindawi; 2021:e8243378 2021. 10.1155/2021/8243378

11. Bonnans, C., J. Chou, and Z. Werb. Remodelling the extracellular matrix in development and disease. Nature Reviews Molecular Cell Biology Nature Publishing Group; 15(12):786–801 2014. 10.1038/nrm3904

12. Cox, T. R., and J. T. Erler. Remodeling and homeostasis of the extracellular matrix: implications for fibrotic diseases and cancer. Disease models & mechanisms The Company of Biologists Ltd; 4(2):165–78 2011. 10.1242/dmm.004077

13. Marozzi, M. et al. Inflammation, Extracellular Matrix Remodeling, and Proteostasis in Tumor Microenvironment. International Journal of Molecular Sciences Multidisciplinary Digital Publishing Institute; 22(15):8102 2021. 10.3390/ijms22158102

14. Mortensen, J. H. et al. The intestinal tissue homeostasis – the role of extracellular matrix remodeling in inflammatory bowel disease. Expert Review of Gastroenterology & Hepatology Taylor & Francis; 13(10):977–993 2019. 10.1080/17474124.2019.1673729

15. Baumgart, D. C., and W. J. Sandborn. Crohn’s disease. The Lancet 380(9853):1590–1605 2012. 10.1016/S0140-6736(12)60026-9

16. Vilardi, A., S. Przyborski, C. Mobbs, A. Rufini, and C. Tufarelli. Current understanding of the interplay between extracellular matrix remodelling and gut permeability in health and disease. Cell Death Discov. Nature Publishing Group; 10(1):258 2024. 10.1038/s41420-024-02015-1

17. Sampaio Moura, N., A. Schledwitz, M. Alizadeh, S. A. Patil, and J.-P. Raufman. Matrix metalloproteinases as biomarkers and therapeutic targets in colitis-associated cancer. Front. Oncol. Frontiers; 13 2024 [cited 2026 Feb 16]. 10.3389/fonc.2023.1325095

18. Biel, C., K. N. Faber, R. A. Bank, and P. Olinga. Matrix metalloproteinases in intestinal fibrosis. J Crohns Colitis 18(3):462–478 2024. 10.1093/ecco-jcc/jjad178

19. Zhihua, L. et al. Fecal Serine Protease Profiling in Inflammatory Bowel Diseases. *Fecal Serine Protease Profiling in Inflammatory Bowel Diseases*. Front. Cell. Infect. Microbiol 10:21 2020. 10.3389/fcimb.2020.00021

20. Vilardi, A., S. Przyborski, C. Mobbs, A. Rufini, and C. Tufarelli. Current understanding of the interplay between extracellular matrix remodelling and gut permeability in health and disease. Cell Death Discov. Nature Publishing Group; 10(1):258 2024. 10.1038/s41420-024-02015-1

21. Glowacki, R. W. P., and E. C. Martens. If You Eat It or Secrete It, They Will Grow: the Expanding List of Nutrients Utilized by Human Gut Bacteria. Journal of Bacteriology American Society for Microbiology; 203(9):10.1128/jb.00481-20 2021. https://doi.org/10.1128/jb.00481-20

22. Aakko, J. et al. A carbohydrate-active enzyme (CAZy) profile links successful metabolic specialization of Prevotella to its abundance in gut microbiota. Sci Rep Nature Publishing Group; 10(1):12411 2020. 10.1038/s41598-020-69241-2

23. Wardman, J. F., R. K. Bains, P. Rahfeld, and S. G. Withers. Carbohydrate-active enzymes (CAZymes) in the gut microbiome. Nat Rev Microbiol Nature Publishing Group; 20(9):542–556 2022. 10.1038/s41579-022-00712-1

24. Galipeau, H. J. et al. Novel Fecal Biomarkers That Precede Clinical Diagnosis of Ulcerative Colitis. Gastroenterology Elsevier BV; 0(0) 2020 [cited 2021 Feb 8]. 10.1053/j.gastro.2020.12.004

25. Annaházi, A. et al. Fecal MMP-9: A New Noninvasive Diperential Diagnostic and Activity Marker in Ulcerative Colitis. 2013 [cited 2021 Nov 11]. 10.1002/ibd.22996

26. Kirov, S. et al. Degradation of the extracellular matrix is part of the pathology of ulcerative colitis. Molecular Omics Royal Society of Chemistry; 15(1):67–76 2019. 10.1039/C8MO00239H

27. Baugh, M. D. et al. Matrix metalloproteinase levels are elevated in inflammatory bowel disease. Gastroenterology Elsevier; 117(4):814–822 1999. 10.1016/S0016-5085(99)70339-2

28. Ribet, D., and P. Cossart. How bacterial pathogens colonize their hosts and invade deeper tissues. Microbes and Infection Elsevier Masson SAS; 17(3):173–183 2015. 10.1016/j.micinf.2015.01.004

29. Singh, B., C. Fleury, F. Jalalvand, and K. Riesbeck. Human pathogens utilize host extracellular matrix proteins laminin and collagen for adhesion and invasion of the host. FEMS Microbiology Reviews 36(6):1122–1180 2012. 10.1111/j.1574-6976.2012.00340.x

30. Hammerschmidt, S., M. Rohde, and K. T. Preissner. Extracellular Matrix Interactions with Gram-Positive Pathogens. Microbiology Spectrum American Society for Microbiology; 7(2):10.1128/microbiolspec.gpp3-0041–2018 2019. 10.1128/microbiolspec.gpp3-0041-2018

31. Porras, A. M. et al. Inflammatory Bowel Disease-Associated Gut Commensals Degrade Components of the Extracellular Matrix. mBio American Society for Microbiology; 13(6):e02201–22 2022. 10.1128/mbio.02201-22

32. Ndeh, D. A. et al. A Bacteroides thetaiotaomicron genetic locus encodes activities consistent with mucin O-glycoprotein processing and N-acetylgalactosamine metabolism. Nat Commun Nature Publishing Group; 16(1):3485 2025. 10.1038/s41467-025-58660-2

33. Wexler, A. G., and A. L. Goodman. An insider’s perspective: Bacteroides as a window into the microbiome. Nat Microbiol Nature Publishing Group; 2(5):17026 2017. 10.1038/nmicrobiol.2017.26

34. Pudlo, N. A. et al. Phenotypic and Genomic Diversification in Complex Carbohydrate-Degrading Human Gut Bacteria. mSystems American Society for Microbiology; 7(1):e00947–21 2022. 10.1128/msystems.00947-21

35. Shin, J. H., G. Tillotson, T. N. MacKenzie, C. A. Warren, H. M. Wexler, and E. J. C. Goldstein. *Bacteroides* and related species: The keystone taxa of the human gut microbiota. Anaerobe 85:102819 2024. 10.1016/j.anaerobe.2024.102819

36. Fansler, R. T., Y. Wu, and W. Zhu. Friend or foe? Contextualizing Bacteroides through the lens of niche remodeling. Trends in Microbiology Elsevier; 0(0) 2025 [cited 2026 Feb 16]. 10.1016/j.tim.2025.11.005

37. Zamani, S., R. Taslimi, A. Sarabi, S. Jasemi, L. A. Sechi, and M. M. Feizabadi. Enterotoxigenic Bacteroides fragilis: A Possible Etiological Candidate for Bacterially-Induced Colorectal Precancerous and Cancerous Lesions. Frontiers in Cellular and Infection Microbiology 9 2020 [cited 2022 Nov 22]. https://www.frontiersin.org/articles/10.3389/fcimb.2019.00449. Accessed: 22 Nov 2022.

38. Cao, Y. et al. Changes in Bacteroides and the microbiota in patients with obstructed colorectal cancer: retrospective cohort study. BJS Open 7(6):zrad105 2023. 10.1093/bjsopen/zrad105

39. Moncrief, J. S. et al. The enterotoxin of Bacteroides fragilis is a metalloprotease. Infection and Immunity American Society for Microbiology; 63(1):175–181 1995. 10.1128/IAI.63.1.175-181.1995

40. Sánchez, E., J. M. Laparra, and Y. Sanz. Discerning the role of bacteroides fragilis in celiac disease pathogenesis. Applied and Environmental Microbiology 78(18):6507–6515 2012. 10.1128/AEM.00563-12/ASSET/6CCA01BC-8BF0-4127-B1E1-42A186B04DE6/ASSETS/GRAPHIC/ZAM9991036290005.JPEG

41. Kawai, K., R. Kamochi, S. Oiki, K. Murata, and W. Hashimoto. Probiotics in human gut microbiota can degrade host glycosaminoglycans. Sci Rep Nature Publishing Group; 8(1):10674 2018. 10.1038/s41598-018-28886-w

42. Hickey, C. A. et al. Colitogenic Bacteroides thetaiotaomicron antigens access host immune cells in a sulfatase-dependent manner via outer membrane vesicles. Cell Host and Microbe Elsevier Inc.; 17(5):672–680 2015. 10.1016/j.chom.2015.04.002

43. Benjdia, A., E. C. Martens, J. I. Gordon, and O. Berteau. Sulfatases and a radical S-adenosyl-L-methionine (AdoMet) enzyme are key for mucosal foraging and fitness of the prominent human gut symbiont, Bacteroides thetaiotaomicron. Journal of Biological Chemistry 286(29):25973–25982 2011. 10.1074/jbc.M111.228841

44. Desai, M. S. et al. A Dietary Fiber-Deprived Gut Microbiota Degrades the Colonic Mucus Barrier and Enhances Pathogen Susceptibility. Cell 167(5):1339–1353.e21 2016. 10.1016/j.cell.2016.10.043

45. Lloyd-Price, J., et al. Multi-omics of the gut microbial ecosystem in inflammatory bowel diseases. Nature Nature Publishing Group; 569(7758):655–662 2019. 10.1038/s41586-019-1237-9

46. Gevers, D. et al. The Treatment-Naive Microbiome in New-Onset Crohn’s Disease. Cell Host & Microbe Elsevier; 15(3):382–392 2014. 10.1016/j.chom.2014.02.005

47. Fang, Z., M. Ma, Y. Wang, W. Dai, Q. Shang, and G. Yu. Degradation and fermentation of hyaluronic acid by *Bacteroides* spp. from the human gut microbiota. Carbohydrate Polymers 334:122074 2024. 10.1016/j.carbpol.2024.122074

48. Wang, Y., M. Ma, W. Dai, Q. Shang, and G. Yu. Bacteroides salyersiae is a potent chondroitin sulfate-degrading species in the human gut microbiota. Microbiome 12(1):41 2024. 10.1186/s40168-024-01768-2

49. Huang, L. et al. dbCAN-seq: a database of carbohydrate-active enzyme (CAZyme) sequence and annotation. Nucleic Acids Research 46(D1):D516–D521 2018. 10.1093/nar/gkx894

50. Zhang, H. et al. dbCAN2: a meta server for automated carbohydrate-active enzyme annotation. Nucleic Acids Research 46(W1):W95–W101 2018. 10.1093/nar/gky418

51. Zheng, J., Q. Ge, Y. Yan, X. Zhang, L. Huang, and Y. Yin. dbCAN3: automated carbohydrate-active enzyme and substrate annotation. Nucleic Acids Research 51(W1):W115–W121 2023. 10.1093/nar/gkad328

52. Rawlings, N. D., A. J. Barrett, P. D. Thomas, X. Huang, A. Bateman, and R. D. Finn. The MEROPS database of proteolytic enzymes, their substrates and inhibitors in 2017 and a comparison with peptidases in the PANTHER database. Nucleic Acids Res 46(D1):D624–D632 2018. 10.1093/nar/gkx1134

53. Naba, A., K. R. Clauser, S. Hoersch, H. Liu, S. A. Carr, and R. O. Hynes. The Matrisome: In Silico Definition and In Vivo Characterization by Proteomics of Normal and Tumor Extracellular Matrices *. Molecular & Cellular Proteomics Elsevier; 11(4) 2012 [cited 2025 Nov 20]. 10.1074/mcp.M111.014647

54. Shao, X. et al. MatrisomeDB 2.0: 2023 updates to the ECM-protein knowledge database. Nucleic Acids Re*s* 51(D1):D1519–D1530 2023. 10.1093/nar/gkac1009

55. Chang, A. et al. BRENDA, the ELIXIR core data resource in 2021: new developments and updates. Nucleic Acids Res 49(D1):D498–D508 2021. 10.1093/nar/gkaa1025

56. Zhou, Y. brendaDb: The BRENDA Enzyme Database. 2025. 10.18129/B9.bioc.brendaDb

57. Dong, J., Y. Cui, and X. Qu. Metabolism mechanism of glycosaminoglycans by the gut microbiota: *Bacteroides* and lactic acid bacteria: A review. Carbohydrate Polymers 332:121905 2024. 10.1016/j.carbpol.2024.121905

58. Wexler, A. G., and A. L. Goodman. An insider’s perspective: Bacteroides as a window into the microbiome. Nat Microbiol Nature Publishing Group; 2(5):1–11 2017. 10.1038/nmicrobiol.2017.26

59. McKee, L. S., S. L. La Rosa, B. Westereng, V. G. Eijsink, P. B. Pope, and J. Larsbrink. Polysaccharide degradation by the Bacteroidetes: mechanisms and nomenclature. Environmental Microbiology Reports 13(5):559–581 2021. 10.1111/1758-2229.12980

60. Martens, E. C. et al. Recognition and Degradation of Plant Cell Wall Polysaccharides by Two Human Gut Symbionts. PLOS Biology Public Library of Science; 9(12):e1001221 2011. 10.1371/journal.pbio.1001221

61. Kaoutari, A. E., F. Armougom, J. I. Gordon, D. Raoult, and B. Henrissat. The abundance and variety of carbohydrate-active enzymes in the human gut microbiota. Nat Rev Microbiol Nature Publishing Group; 11(7):497–504 2013. 10.1038/nrmicro3050

62. Xu, J. et al. A Genomic View of the Human-Bacteroides thetaiotaomicron Symbiosis. Science American Association for the Advancement of Science; 299(5615):2074–2076 2003. 10.1126/science.1080029

63. Sonnenburg, E. D. et al. Specificity of Polysaccharide Use in Intestinal Bacteroides Species Determines Diet-Induced Microbiota Alterations. Cell Elsevier; 141(7):1241–1252 2010. 10.1016/j.cell.2010.05.005

64. Xu, J. et al. Evolution of Symbiotic Bacteria in the Distal Human Intestine. PLOS Biology Public Library of Science; 5(7):e156 2007. 10.1371/journal.pbio.0050156

65. Hehemann, J.-H., G. Correc, T. Barbeyron, W. Helbert, M. Czjzek, and G. Michel. Transfer of carbohydrate-active enzymes from marine bacteria to Japanese gut microbiota. 2010 [cited 2018 Aug 1]. 10.1038/nature08937

66. Raghavan, V., and E. A. Groisman. Species-Specific Dynamic Responses of Gut Bacteria to a Mammalian Glycan. Journal of Bacteriology American Society for Microbiology; 197(9):1538–1548 2015. 10.1128/jb.00010-15

67. Ulmer, J. E. et al. Characterization of Glycosaminoglycan (GAG) Sulfatases from the Human Gut Symbiont *Bacteroides thetaiotaomicron* Reveals the First GAG-specific Bacterial Endosulfatase*. Journal of Biological Chemistry 289(35):24289–24303 2014. 10.1074/jbc.M114.573303

68. Overbeeke, A., B. Hausmann, G. Nikolov, F. C. Pereira, C. W. Herbold, and D. Berry. Nutrient niche specificity for glycosaminoglycans is reflected in polysaccharide utilization locus architecture of gut Bacteroides species. Front. Microbiol. Frontiers; 13 2022 [cited 2025 Dec 9]. 10.3389/fmicb.2022.1033355

69. Hyun, Y.-J., I.-H. Jung, and D.-H. Kim. Expression of heparinase I of *Bacteroides stercoris* HJ-15 and its degradation tendency toward heparin-like glycosaminoglycans. Carbohydrate Research 359:37–43 2012. 10.1016/j.carres.2012.05.023

70. Houston, S., G. W. Blakely, A. McDowell, L. Martin, and S. Patrick. Binding and degradation of fibrinogen by Bacteroides fragilis and characterization of a 54 kDa fibrinogen-binding protein. Microbiology Microbiology Society,; 156(8):2516–2526 2010. 10.1099/mic.0.038588-0

71. Chen, Y., T. Kinouchi, K. Kataoka, S. Akimoto, and Y. Ohnishi. Purification and Characterization of a Fibrinogen-Degrading Protease in Bacteroides fragilis Strain YCH46. Microbiology and Immunology 39(12):967–977 1995. 10.1111/j.1348-0421.1995.tb03300.x

72. Gibson, S. A. W., and G. T. Macfarlane. Characterization of Proteases Formed by Bacteroides fragilis. Microbiology Microbiology Society,; 134(8):2231–2240 1988. 10.1099/00221287-134-8-2231

73. Stepen, E. K., and D. J. Hentges. Hydrolytic enzymes of anaerobic bacteria isolated from human infections. Journal of Clinical Microbiology American Society for Microbiology; 14(2):153–156 1981. 10.1128/jcm.14.2.153-156.1981

74. Wikström, M., and A. Linde. Ability of oral bacteria to degrade fibronectin. Infection and Immunity American Society for Microbiology; 51(2):707–711 1986. 10.1128/iai.51.2.707-711.1986

75. Rudek, W., and R. U. Haque. Extracellular enzymes of the genus Bacteroides. Journal of Clinical Microbiology American Society for Microbiology; 4(5):458–460 1976. 10.1128/jcm.4.5.458-460.1976

76. Griesemer, M., J. A. Kimbrel, C. E. Zhou, A. Navid, and P. D’haeseleer. Combining multiple functional annotation tools increases coverage of metabolic annotation. BMC Genomics 19(1):948 2018. 10.1186/s12864-018-5221-9

77. Rembeza, E., and M. K. M. Engqvist. Experimental and computational investigation of enzyme functional annotations uncovers misannotation in the EC 1.1.3.15 enzyme class. PLOS Computational Biology Public Library of Science; 17(9):e1009446 2021. 10.1371/journal.pcbi.1009446

78. McDonald, A. G., and K. F. Tipton. Enzyme nomenclature and classification: the state of the art. The FEBS Journal 290(9):2214–2231 2023. 10.1111/febs.16274

79. Owen, C. A. Proteinases and Oxidants as Targets in the Treatment of Chronic Obstructive Pulmonary Disease. Proc Am Thorac Soc 2(4):373–385 2005. 10.1513/pats.200504-029SR

80. Serwanja, J., A. C. Wieland, A. Haubenhofer, H. Brandstetter, and E. Schönauer. A conserved strategy to attack collagen: The activator domain in bacterial collagenases unwinds triple-helical collagen. Proceedings of the National Academy of Sciences Proceedings of the National Academy of Sciences; 121(16):e2321002121 2024. 10.1073/pnas.2321002121

81. Jones, E. J. et al. The Uptake, Trapicking, and Biodistribution of Bacteroides thetaiotaomicron Generated Outer Membrane Vesicles. Front. Microbiol. Frontiers; 11 2020 [cited 2026 Mar 4]. 10.3389/fmicb.2020.00057

82. Michielan, A., and R. D’Incà. Intestinal Permeability in Inflammatory Bowel Disease: Pathogenesis, Clinical Evaluation, and Therapy of Leaky Gut. Mediators of Inflammation Hindawi; 2015:e628157 2015. 10.1155/2015/628157

83. Yu, L. C.-H. Microbiota dysbiosis and barrier dysfunction in inflammatory bowel disease and colorectal cancers: exploring a common ground hypothesis. Journal of Biomedical Science BioMed Central; 25(1):79 2018. 10.1186/s12929-018-0483-8

84. Turpin, W. et al. Increased Intestinal Permeability Is Associated With Later Development of Crohn’s Disease. Gastroenterology 159(6):2092–2100.e5 2020. 10.1053/j.gastro.2020.08.005

85. Vrakas, S. et al. Intestinal Bacteria Composition and Translocation of Bacteria in Inflammatory Bowel Disease. PLOS ONE Public Library of Science; 12(1):e0170034 2017. 10.1371/journal.pone.0170034

86. Kleessen, B., A. J. Kroesen, H. J. Buhr, and M. Blaut. Mucosal and Invading Bacteria in Patients with Inflammatory Bowel Disease Compared with Controls. Scandinavian Journal of Gastroenterology Taylor & Francis; 37(9):1034–1041 2002. 10.1080/003655202320378220

87. Kessler, S., H. Rho, G. West, C. Fiocchi, J. Drazba, and C. De La Motte. Hyaluronan (HA) Deposition Precedes and Promotes Leukocyte Recruitment in Intestinal Inflammation. Clinical Translational Sci 1(1):57–61 2008. 10.1111/j.1752-8062.2008.00025.x

88. Termeer, C. et al. Oligosaccharides of Hyaluronan activate dendritic cells via toll-like receptor 4. The Journal of experimental medicine Rockefeller University Press; 195(1):99–111 2002. 10.1084/jem.20001858

89. Houghton, A. M. et al. Elastin fragments drive disease progression in a murine model of emphysema. J Clin Invest American Society for Clinical Investigation; 116(3):753–759 2006. 10.1172/JCI25617

90. Pfister, R. R., J. L. Haddox, C. I. Sommers, and K. W. Lam. Identification and synthesis of chemotactic tripeptides from alkali-degraded whole cornea. A study of N-acetyl-proline-glycine-proline and N-methyl-proline-glycine-proline. Investigative Ophthalmology & Visual Science 36(7):1306–1316 1995.

91. Galipeau, H. J., A. Caminero, and E. F. Verdu. Increased Bacterial Proteolytic Activity Detected Before Diagnosis of Ulcerative Colitis. Inflammatory Bowel Diseases 27(12):e144 2021. 10.1093/ibd/izab144

92. Mills, R. H. et al. Multi-omics analyses of the ulcerative colitis gut microbiome link Bacteroides vulgatus proteases with disease severity. Nat Microbiol Nature Publishing Group; 7(2):262–276 2022. 10.1038/s41564-021-01050-3

93. Caminero, A., M. Guzman, J. Libertucci, and A. E. Lomax. The emerging roles of bacterial proteases in intestinal diseases. Gut Microbes Taylor & Francis; 15(1):2181922 2023. 10.1080/19490976.2023.2181922

94. Rondeau, L. E. et al. Proteolytic bacteria expansion during colitis amplifies inflammation through cleavage of the external domain of PAR2. Gut Microbes Taylor & Francis; 16(1):2387857 2024. 10.1080/19490976.2024.2387857

95. Jørgensen, A. B., I. Jonsson, L. Friis-Hansen, and B. Brandstrup. Collagenase-producing bacteria are common in anastomotic leakage after colorectal surgery: a systematic review. Int J Colorectal Dis 38(1):275 2023. 10.1007/s00384-023-04562-y

96. The UniProt Consortium. UniProt: the Universal Protein Knowledgebase in 2025. Nucleic Acids Res 53(D1):D609–D617 2025. 10.1093/nar/gkae1010

97. Ahmad, S. et al. The UniProt website API: facilitating programmatic access to protein knowledge. Nucleic Acids Res 53(W1):W547–W553 2025. 10.1093/nar/gkaf394

98. Seemann, T. Prokka: rapid prokaryotic genome annotation. Bioinformatics 30(14):2068–2069 2014. 10.1093/bioinformatics/btu153

99. Huerta-Cepas, J. et al. eggNOG 5.0: a hierarchical, functionally and phylogenetically annotated orthology resource based on 5090 organisms and 2502 viruses. Nucleic Acids Research 47(D1):D309–D314 2019. 10.1093/nar/gky1085

100. Cantalapiedra, C. P., A. Hernández-Plaza, I. Letunic, P. Bork, and J. Huerta-Cepas. eggNOG-mapper v2: Functional Annotation, Orthology Assignments, and Domain Prediction at the Metagenomic Scale. Molecular Biology and Evolution 38(12):5825–5829 2021. 10.1093/molbev/msab293

101. Buchfink, B., K. Reuter, and H.-G. Drost. Sensitive protein alignments at tree-of-life scale using DIAMOND. Nat Methods Nature Publishing Group; 18(4):366–368 2021. 10.1038/s41592-021-01101-x

102. Drula, E., M.-L. Garron, S. Dogan, V. Lombard, B. Henrissat, and N. Terrapon. The carbohydrate-active enzyme database: functions and literature. Nucleic Acids Res 50(D1):D571–D577 2022. 10.1093/nar/gkab1045

103. Larralde, M., and G. Zeller. PyHMMER: a Python library binding to HMMER for epicient sequence analysis. Bioinformatics 39(5):btad214 2023. 10.1093/bioinformatics/btad214

104. Finn, R. D., J. Clements, and S. R. Eddy. HMMER web server: interactive sequence similarity searching. Nucleic Acids Res 39(suppl_2):W29–W37 2011. 10.1093/nar/gkr367

105. Caporaso, J. G. et al. Global patterns of 16S rRNA diversity at a depth of millions of sequences per sample. Proceedings of the National Academy of Sciences Proceedings of the National Academy of Sciences; 108(supplement_1):4516–4522 2011. 10.1073/pnas.1000080107

106. Ladner, Y. D., M. Alini, and A. R. Armiento. The Dimethylmethylene Blue Assay (DMMB) for the Quantification of Sulfated Glycosaminoglycans. In: M. J. Stoddart, E. Della Bella, and A. R. Armiento, editors. Cartilage Tissue Engineering. New York, NY: Springer US; 2023 [cited 2025 Aug 1]. pp. 115–121. 10.1007/978-1-0716-2839-3_9

107. Frank E Harrel Jr. Hmisc: Harrell Miscellaneous. 2025. 10.32614/CRAN.package.Hmisc

108. Oksanen J et al. vegan: Community Ecology Package. 2025. 10.32614/CRAN.package.vegan

109. Posit team. RStudio: Integrated Development Environment for R. Boston, MA: Posit Software, PBC; 2025.

