## Supplemental File for "Comparative Genomic and Functional Profiling of ECM-Targeting Enzymes in *Bacteroides*, a Key Genus of the Human Gut Microbiome"

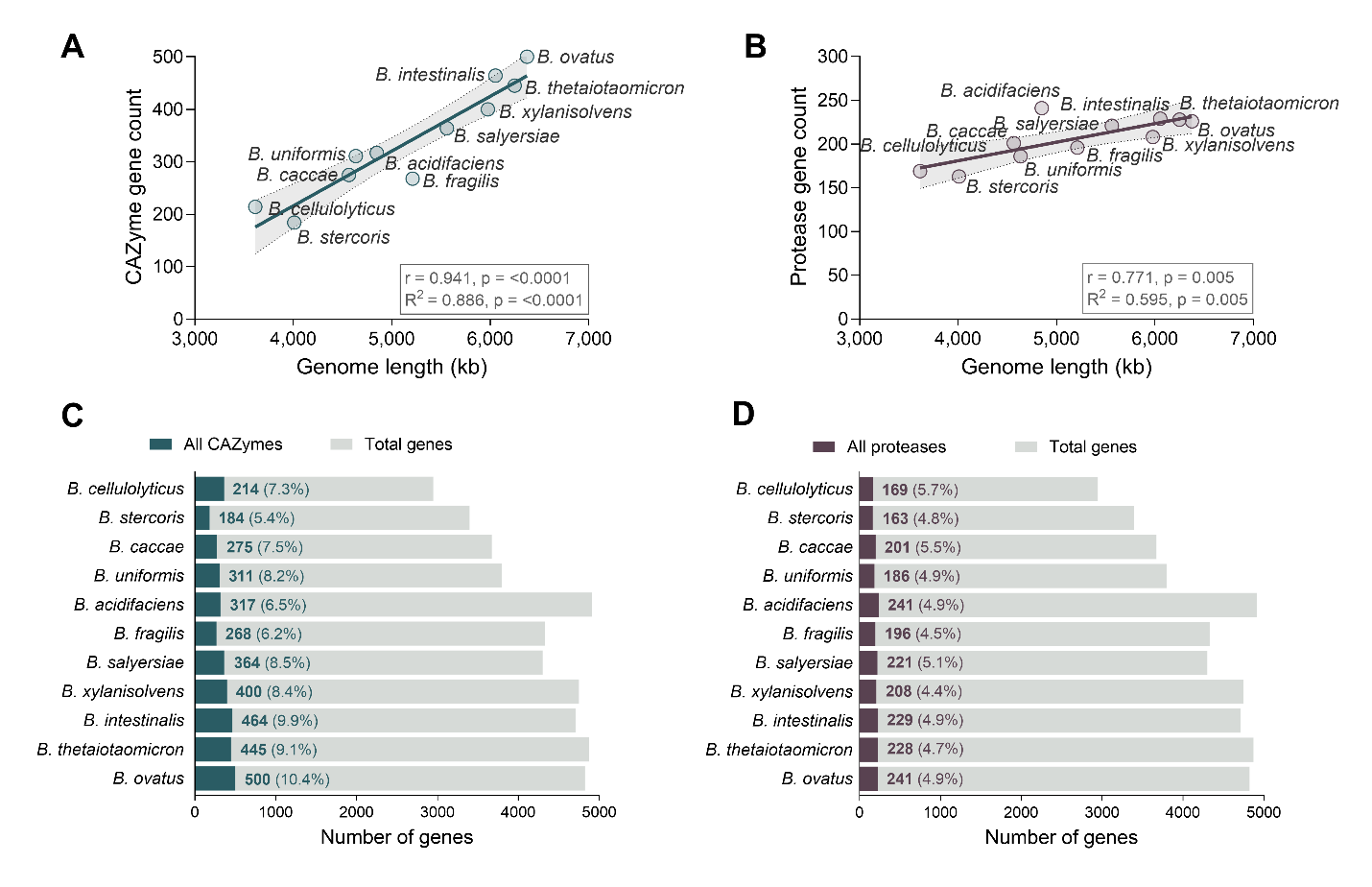

**Supplementary Figure 1**. **Abundance of total CAZyme and protease genes identified across *Bacteroides* genomes**. **(A-B)** Relationship between genome length and the total number of genes identified as (A) CAZymes or (B) proteases. Each species is denoted by a point, and the line represents a linear regression with 95% confidence intervals. The Pearson correlation coefficient (r), the coefficient of determination (R²), and their associated p value are displayed. **(C)** Total CAZymes and **(D)** total proteases as a percentage of the total encoded genes in each species’ genome. Species are ordered by ascending genome length.

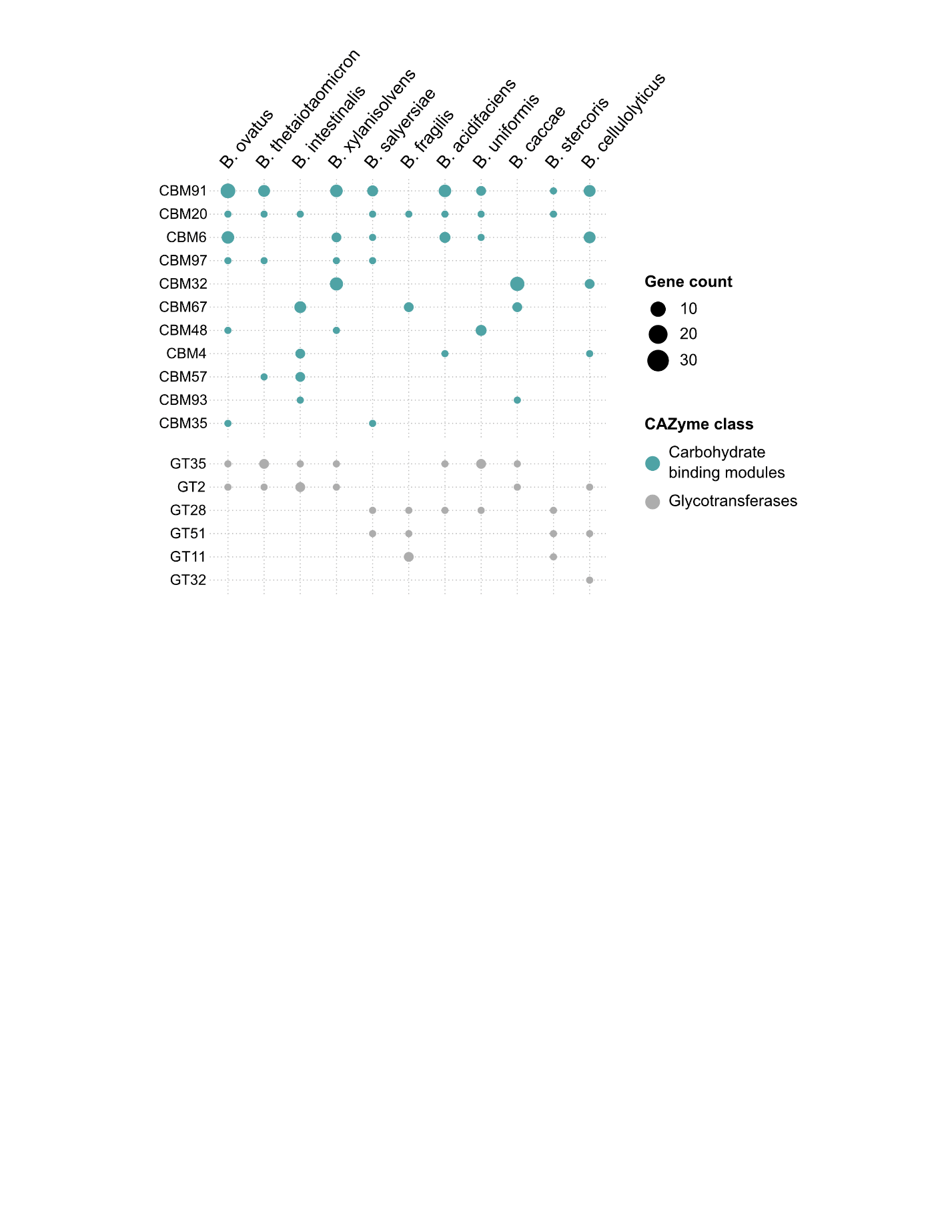

**Supplementary Figure 2. Family level gene copy number of ECM-associated CAZymes belonging to the CBM and GT classes.** Dotplot of gene copy numbers for each family in the carbohydrate binding modules and glycotransferase classes per species. Species are ordered as columns by descending genome length. Families within each class are ordered first by decreasing species prevalence and then by decreasing gene copy number. Dot size corresponds to the gene copy number in each family per species.

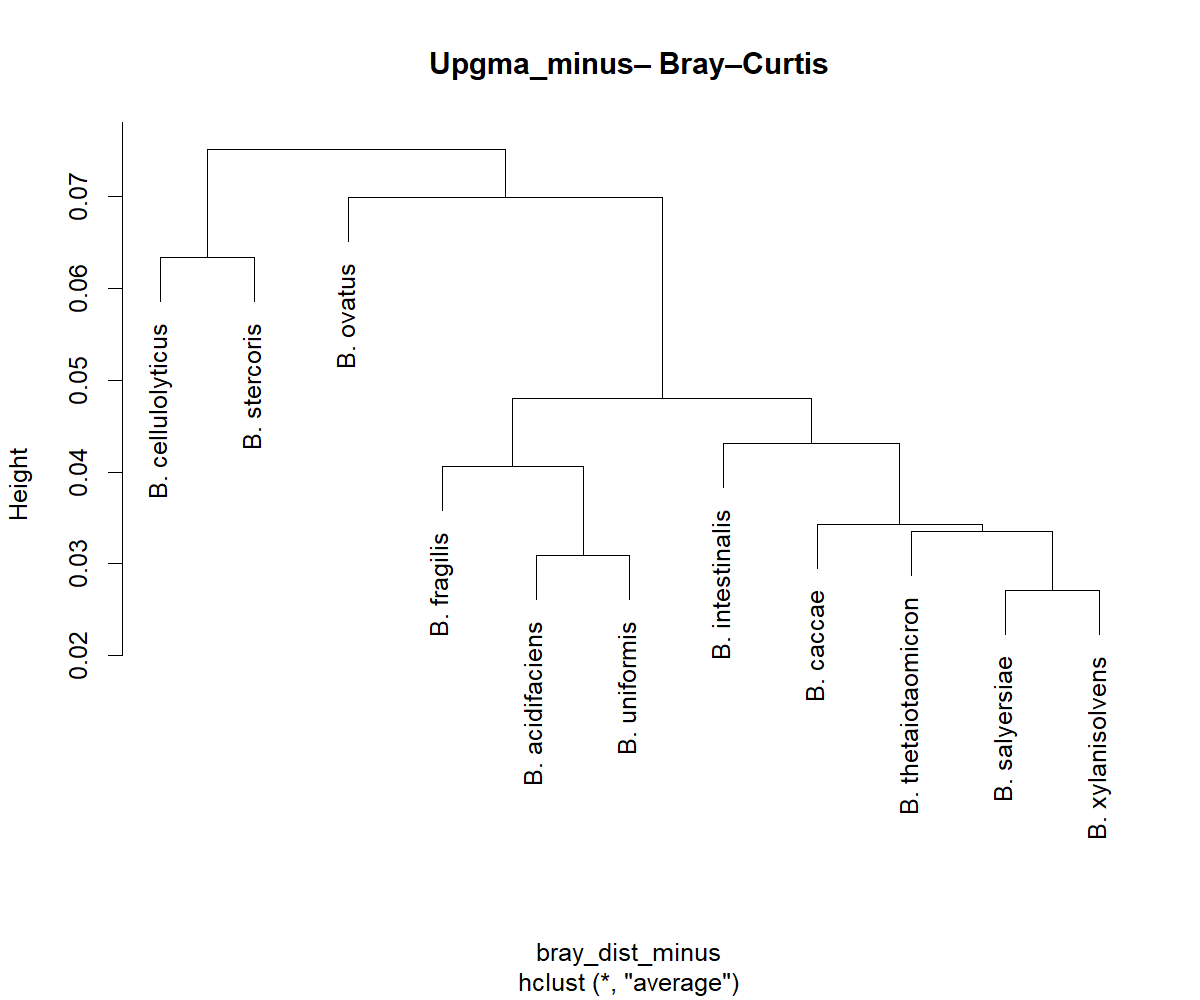

**Supplementary Figure 3. Hierarchical clustering dendrogram of species based on Bray-Curtis dissimilarity.** Clustering was calculated from substrate-level relative gene count profiles per species using the average linkage (UPGMA) algorithm. Branch lengths represent the degree of dissimilarity between species’ substrate-level gene profiles.

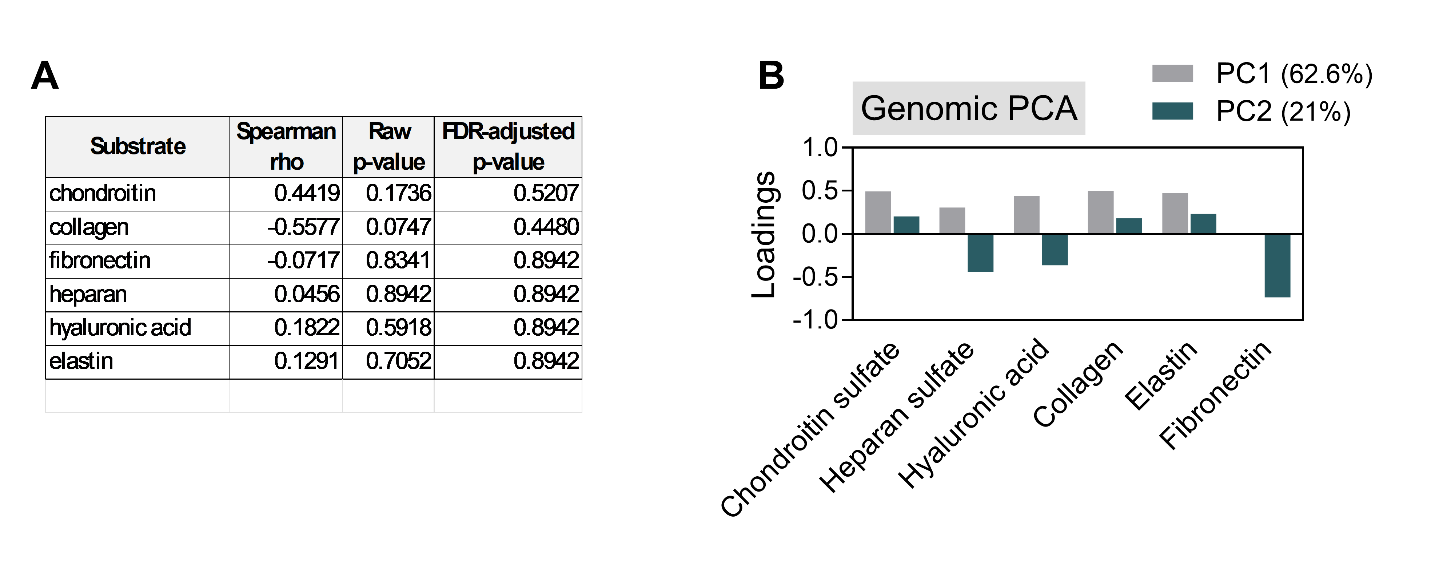

**Supplementary Figure 4. Correlation analysis and principal component loadings of substrate-level gene profiles.** **(A)** Spearman’s rank correlation coefficients between predicted genomic gene counts and experimental activity across six substrates. Corresponding raw p value and false discovery rate (FDR)-adjusted p value are shown. **(B)** Principal component analysis (PCA) loadings per substrate for PC1 and PC2 derived from genomic prediction PCA analysis. The percentage variance explained by each component is indicated in the legend.

**Supplementary Table 1. Core human ECM terms used to build ECM enzyme database listed by separate functional classifications.**

| **Functional Class** | **Extracellular Matrix Terms** |
| --- | --- |
| Proteins | collagen, elastin, fibrillin, matrix protein, chondroadherin, discoidin, otogelin, otolin, periostin, tectorin, tenascin, thrombospondin |
| Proteoglycans | aggrecan, biglycan, brevican, decorin, keratocan, perlecan, serglycin, versican |
| Glycoproteins | agrin, ameloblastin, amelogenin, asporin, dermatopontin, extracellular phosphoglycoprotein, fibronectin, fibrinogen, fibromodulin, fibulin, gliomedin, hemicentin, hevin, kielin, laminin, matrilin, multimerin, nephrocan, nephronectin, neurocan, nidogen, podocan, opticin, osteoglycin, osteomodulin, osteonectin, sialoprotein, vitrin, vitronectin |
| Glycosaminoglycans | chondroitin, hyaluron, hyaluronic acid, dermatan, heparan, keratan |
| Cell adhesion molecules | focal adhesion, integrin, netrin, paxillin, talin, vinculin, zyxin |
| Other ECM components | dentin, extracellular matrix, matrix phosphoglycoprotein |

**Supplementary Table 2. Intestinal ECM terms listed by separate functional classifications.**

| **Functional Class** | **Extracellular Matrix Terms** |
| --- | --- |
| Proteins | collagen, elastin, laminin, osteonectin |
| Proteoglycans | agrin, biglycan, decorin, fibromodulin, perlecan, serglycin, versican |
| Glycoproteins | fibronectin, fibrillin, fibrinogen, fibulin, laminin, nidogen, periostin, tenascin, thrombospondin, vitronectin |
| Glycosaminoglycans | chondroitin, hyaluron, hyaluronic acid, dermatan, heparan, keratan |
